# Proximity-based activation of AURORA A by MPS1 potentiates error correction

**DOI:** 10.1101/2024.06.11.598300

**Authors:** Nelson Leça, Francisca Barbosa, Sergi Rodriguez-Calado, Margarida Moura, Paulo D. Pedroso, Inês Pinto, Arianna E. Verza, Tanja Bange, Claudio E. Sunkel, Marin Barisic, Thomas J. Maresca, Carlos Conde

## Abstract

Faithfull cell division relies on mitotic chromosomes becoming bioriented with each pair of sister kinetochores bound to microtubules oriented toward opposing spindle poles. Erroneous kinetochore-microtubule attachments often form during early mitosis, but are destabilized through the phosphorylation of outer kinetochore proteins by centromeric AURORA B kinase (ABK) and centrosomal AURORA A kinase (AAK), thus allowing for re-establishment of attachments until biorientation is achieved. MPS1-mediated phosphorylation of NDC80 has also been shown to directly weaken the kinetochore-microtubule interface in yeast. In human cells, MPS1 has been proposed to transiently accumulate at end-on attached kinetochores and phosphorylate SKA3 to promote microtubule release. Whether MPS1 directly targets NDC80 and/or promotes the activity of AURORA kinases in metazoans remains unclear. Here, we report a novel mechanism involving communication between kinetochores and centrosomes, wherein MPS1 acts upstream of AAK to promote error correction. MPS1 on pole-proximal kinetochores phosphorylates the C-lobe of AAK thereby increasing its activation at centrosomes. This proximity-based activation ensures the establishment of a robust AAK activity gradient that locally destabilizes mal-oriented kinetochores near spindle poles. Accordingly, MPS1 depletion from *Drosophila* cells causes severe chromosome misalignment and erroneous kinetochore-microtubule attachments, which can be rescued by tethering either MPS1 or constitutively active AAK mutants to centrosomes. Proximity-based activation of AAK by MPS1 also occurs in human cells to promote AAK-mediated phosphorylation of the NDC80 N-terminal tail. These findings uncover an MPS1-AAK cross-talk that is required for efficient error correction, showcasing the ability of kinetochores to modulate centrosome outputs to ensure proper chromosome segregation.

## RESULTS

While it has long been appreciated that MPS1 contributes to error correction in human cells, defining whether it plays a direct or indirect role in the process has proven difficult. Initial evidence suggested that MPS1 could contribute to error correction indirectly via activation of centromeric AURORA B kinase (ABK) [1–3], however, other studies concluded that MPS1 acted downstream of ABK [4] and reported that MPS1 inhibition did not reduce ABK’s ability to phosphorylate known substrates in cells [5,6]. A direct role for kinetochore-associated MPS1 in error correction was later supported by the identification of kinetochore-microtubule attachment factors as MPS1 substrates in human and budding yeast. In humans, a pool of MPS1 localizes to end-on attached kinetochores prior to error correction [7] and MPS1-mediated phosphorylation of the SKA complex, which accumulates at end-on attachments to stabilize them, hinders its association with dynamic microtubules *in vitro* and is required for error correction in cells [8]. Similarly, in budding yeast, phosphorylation of the attachment factor NDC80 by kinetochore-associated MPS1 weakens the binding of kinetochore particles to microtubules *in vitro* and contributes to error correction and accurate chromosome segregation *in vivo* [9]. Thus, efficient error correction in yeast and humans relies on direct inputs from both MPS1 and AURORA kinases via phosphorylation of kinetochore-microtubule attachment factors. However, whether MPS1 directly targets NDC80 and/or promotes the activity of AURORA kinase in metazoans are open questions.

*Drosophila melanogaster* offers a unique model to further dissect error correction because key players in the pathway (MPS1, AURORA kinases and the NDC80 complex) are conserved but kinetochore composition in flies is divergent relative to human and budding yeast in notable ways. For example, flies do not have a functional SKA complex but, like humans, possess centrosome-enriched AURORA A kinase (AAK) and centromere/kinetochore-localized ABK while budding yeast has only a single AURORA kinase orthologue (Ipl1). AURORA kinases play a conserved role in error correction by targeting NDC80, but the localization of AAK and ABK imparts a spatial component to error correction that is likely absent in budding yeast. Indeed, AAK contributes to a polar error correction pathway in flies and mammals via phosphorylation of NDC80 at pole-proximal kinetochores [10,11]. In flies, MPS1 also localizes to microtubule-attached kinetochores [12], but if it contributes to error correction then it does so in the absence of the SKA complex and in the presence of spatial cues from AURORA kinases.

### Polar chromosomes promote AAK activation at centrosomes in an MPS1-dependent manner

The activation status of AURORA kinases throughout the cell cycle was assessed in *D. melanogaster* S2 cells using antibodies against the phosphorylated T-loops of ABK (ABK^T209Ph^) and AAK (AAK^T311Ph^), the latter of which was distinguished from cross-reactive ABK^T209Ph^ by its colocalization with the *Drosophila* Pericentrin-Like Protein (DPLP). Active AAK^T311Ph^ was undetectable at interphase centrosomes but increased significantly during mitosis peaking in prophase and declining from prometaphase through metaphase and anaphase (Figures 1A and 1B). Mis-aligned and unattached kinetochores stimulated centrosomal AAK activation since MG132-treated cells in prometaphase or co-treated with colchicine to depolymerize microtubules exhibited comparable levels of AAK^T311Ph^ that were significantly higher than AAK^T311Ph^ levels in metaphase-arrested cells (Figure S1A and S1B). Interestingly, robust activation of AAK required MPS1 activity as AAK^T311Ph^ levels at centrosomes remained low throughout mitosis in cells depleted of MPS1 relative to controls (Figures 1A and 1B).

**FIGURE 1.**
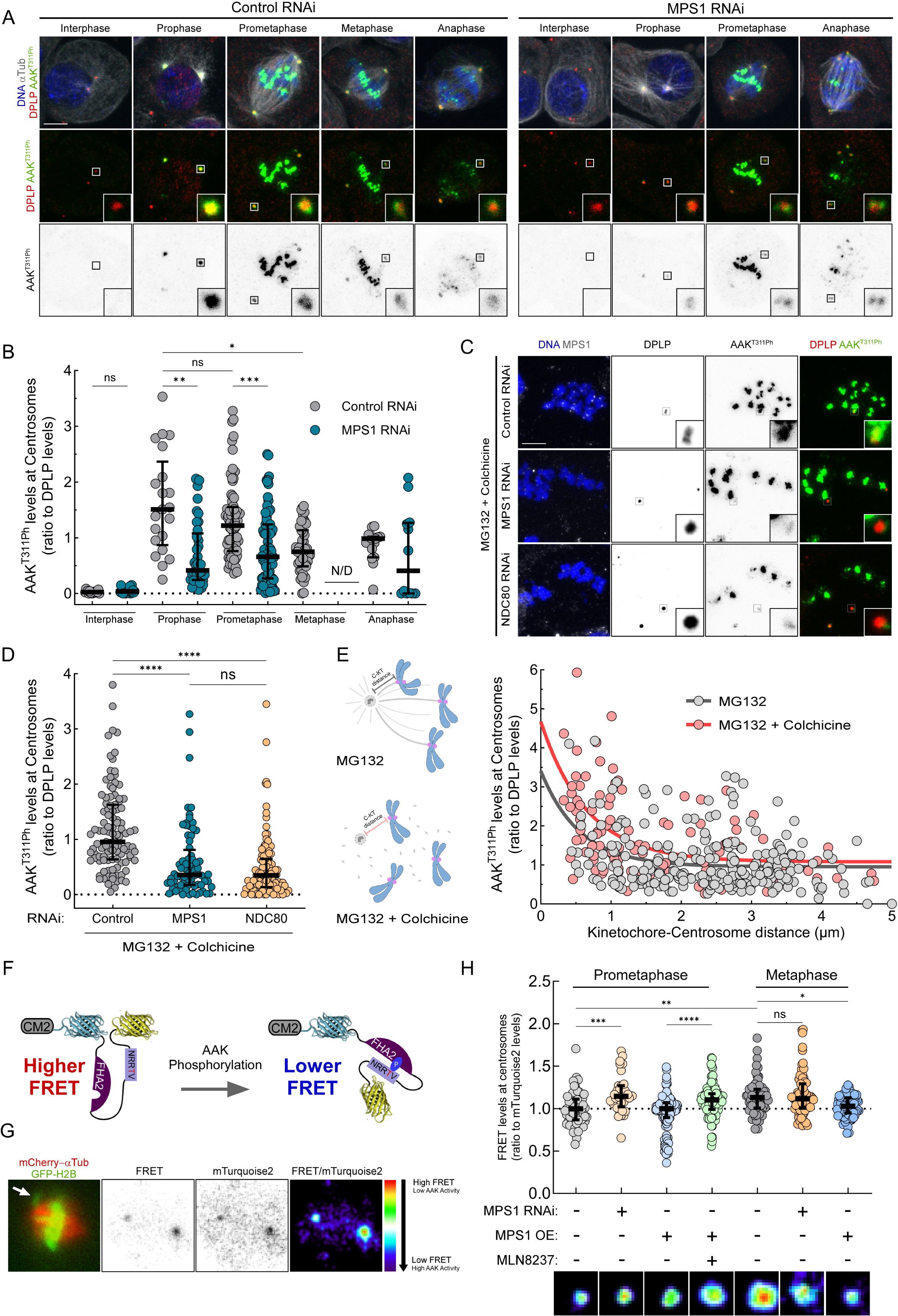
Polar chromosomes promote AAK activation at centrosomes in an MPS1-dependent manner. (A) Representative immunofluorescence images of AAK T311 phosphorylation (AAK^T311Ph^) in control and MPS1-depleted *Drosophila* S2 cells in the indicated cell cycle stages. Insets depict magnifications of selected centrosomes. (B) Quantification of the levels of AAK T311 phosphorylation (AAK^T311Ph^) relative to DPLP at centrosomes of control and MPS1-depleted *Drosophila* S2 cells in the indicated cell cycle stages, as presented in (A). All values were normalized to the mean value determined for prometaphase control cells, which was set to 1. (C) Representative immunofluorescence images of AAK T311 phosphorylation (AAK^T311Ph^) in control, MPS1-depleted or NDC80-depleted *Drosophila* S2 cells treated with 20 µM MG132 for 3 h and 30µM colchicine for 2h. Insets depict magnifications of selected centrosomes. (D) Quantification of the levels of AAK T311 phosphorylation (AAK^T311Ph^) relative to DPLP at centrosomes of control, MPS1-depleted and NDC80-depleted *Drosophila* S2 cells treated with 20 µM MG132 for 3 h and 30 µM colchicine for 2h in the indicated conditions, as presented in (C). All values were normalized to the mean value determined for control cells, which was set to 1. (E) Scatter plot dispersion of fluorescence intensity levels of AAK T311 phosphorylation (AAK^T311Ph^) relative to DPLP at each centrosome vs the 3D distance between that centrosome and the closest kinetochore (measured as indicated in the schematic representation at the left) of control *Drosophila* S2 cells treated with 20 µM MG132 for 3 h or cells treated with and 20 µM MG132 for 3 h and 30 µM colchicine for 2h. All values were normalized to the mean value determined for control cells treated with 20 µM MG132 for 3 h, which was set to 1. The two lines depicted represent a one-decay exponential fit for the indicated experimental conditions. Equations of the depicted exponential fits: y=2,46*e^-1,80x^+0.95 (grey); y=3,60*e^-1,52x^+1.08 (red). (F) Schematic of the centrosome-targeted CM2-AURORA FRET sensor used in (G,H). (G) Representative images of the FRET reporter in a control cell with one polar chromosome (white arrow) near the left centrosome. The FRET emission ratio images “FRET/mTurquoise2” are pseudo-colored, according to the color wedge in display. All images are from a single selected Z-slice in the indicated channels. (H) Quantification of the levels of FRET emission relative to total mTurquoise2 levels at centrosomes of cells and representative FRET emission ratio images “FRET/mTurquoise2” pseudo-colored in the indicated conditions. All images are from a single selected Z-slice and depict magnifications of selected centrosomes from the indicated conditions. All values were normalized to the mean value determined for prometaphase control cells, which was set to 1. Data information: Data in (B), (D) and (H) are presented as median ± interquartile range. Statistical analysis was performed using a Kruskal–Wallis test for multiple comparisons. P values: ns, not significant; *< 0.05; **< 0.01; ***< 0.001; ****< 0.0001. Scale bar: 5 μm. See also Figure S1.

The effect of MPS1 depletion on AAK activation was next examined following depolymerization of microtubules with colchicine since MPS1 becomes enriched at unattached kinetochores via direct interaction with the NDC80 complex [13–18]. High levels of AAK^T311Ph^ were observed in control cells, but AAK activation was significantly compromised following depletion of either MPS1 or NDC80, which mis-localized MPS1 from kinetochores (Figures 1C, 1D and S1C). Thus, kinetochore-associated MPS1 at mis-aligned and unattached kinetochores was required for the efficient activation of centrosomal AAK in mitosis. It was noteworthy that the effect of inhibiting MPS1 was specific to AAK because ABK^T209Ph^ levels at centromeres remained high throughout mitosis or following colchicine treatment in MPS1- and NDC80-depleted cells relative to control RNAi-treated cells (Figures S1D-S1G). Centrosomal AAK^T311Ph^ levels were plotted as a function of distance to the nearest kinetochore to assess whether there was a spatial component to the MPS1-dependent activation of AAK. There was a clear correlation between the extent of AAK activation at centrosomes and proximity to kinetochores such that higher levels of centrosomal AAK^T311Ph^ were measured when kinetochores were present within a ∼1 μm radius of the centrosome (Figure 1E). While the spatial effect was observed in both colchicine and untreated cells, the extent of kinetochore-proximal AAK activation was generally higher in colchicine treated cells consistent with there being elevated levels of MPS1 at unattached kinetochores.

A FRET-based biosensor for AURORA kinase activity [11,19] was next used to study if MPS1-dependent activation of centrosomal AAK affected phosphorylation of downstream substrates. The reporter, which undergoes a conformational change upon phosphorylation that results in reduced FRET efficiency [20], was tethered to centrosomes via an N-terminal fusion to the CM2 domain of Centrosomin [21] (Figure 1F). Centrosome targeting put the reporter in close proximity to AAK, but higher phosphorylation (lower FRET) of the reporter was only observed at centrosomes with nearby polar chromosomes (Figure 1G). Quantification of FRET emission ratios in prometaphase revealed a significant decrease in phosphorylation (higher FRET) of the reporter following MPS1 depletion (Figure 1H). FRET emission ratios were comparable in control and MPS1 over-expressing (MPS1 OE) prometaphase cells suggesting that the centrosome-targeted sensor approaches maximal phosphorylation in prometaphase. Phosphorylation of the reporter in MPS1 over-expressing prometaphase cells required AAK activity since treatment with the inhibitor MLN8237 at a concentration previously determined to specifically inhibit AAK in *Drosophila* S2 cells [11] decreased phosphorylation of the reporter to the same extent as that measured in MPS1-depleted cells (Figure 1H). Consistent with the prior observation of decreased AAK activation at centrosomes in metaphase (Figures 1A and 1B), the CM2-fused reporter exhibited reduced phosphorylation at metaphase centrosomes of control and MPS1-depleted cells (Figure 1H). However, over-expression of MPS1 increased phosphorylation of the reporter in metaphase to comparable levels as that measured in prometaphase cells (Figure 1H). Taken together these findings support the conclusion that the proximity-based activation of AAK at centrosomes by MPS1 potentiates the phosphorylation of downstream AAK substrates.

### MPS1-mediated phosphorylation of AAK C-lobe potentiates AAK auto-activation

*In vitro* kinase assays with purified human orthologues of His-AAK and GST-MPS1 were employed to gain mechanistic insights into how MPS1 activates AAK. Co-incubation of AAK with MPS1 resulted in a ∼4-fold increase in the phosphorylation of AAK activation loop (AAK^T288Ph^, equivalent to *Drosophila* AAK^T311Ph^), which was reduced by half in the presence of the MPS1 inhibitor Cpd-5 [22] (Figures 2A and 2B). The AAK inhibitor MLN8237 prevented MPS1-induced AAK activation indicating that MPS1 does not directly phosphorylate the T-loop (T288) but rather stimulates its auto-phosphorylation. Mass spectrometry of *in vitro* phosphorylated AAK identified a putative MPS1 target site located at a conserved residue (T337/T360 in human and *Drosophila* AAK respectively) that is predicted by AlphaFold3 to be positioned in the C-lobe αG-helix with close proximity to the T-loop (Figure S2A and S2B).

**FIGURE 2.**
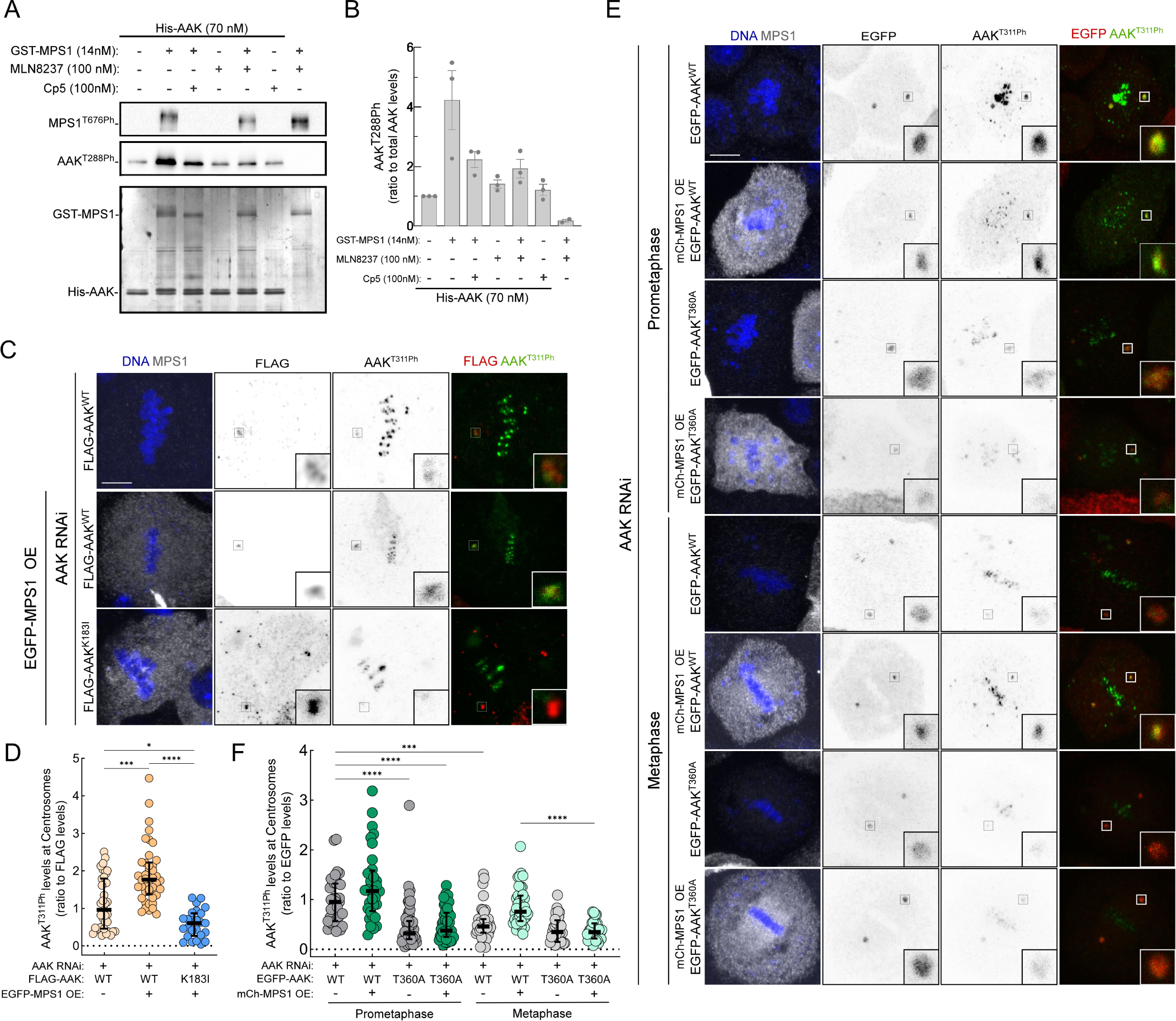
MPS1-mediated phosphorylation of AAK C-lobe potentiates AAK auto-activation. (A) *In vitro* kinase assay with purified human orthologues of AAK and MPS1. Western blot analysis of human AAK T288 activating T-loop autophosphorylation (AAK^T288Ph^) and human MPS1 T676 activating T-loop autophosphorylation (MPS1^T676Ph^). Recombinant His6-AAK was incubated with 100μM ATP during 30 min at 30°C in the indicated conditions. Total protein levels were assessed by gel silver staining. (B) Quantification of human AAK T288 activating T-loop phosphorylation (AAK^T288Ph^) blot intensity relative to total AAK levels in the indicated conditions, as presented in (A). All values were normalized to the mean value determined for basal AAK T288 phosphorylation, which was set to 1. (C) Representative immunofluorescence images of AAK T311 phosphorylation (AAK^T311Ph^) in metaphase *Drosophila* S2 cells depleted of endogenous AAK and expressing FLAG-AAK^WT^ or FLAG-AAK^K183I^ in the indicated conditions. Insets depict magnifications of selected centrosomes. (D) Quantification of the levels of AAK T311 phosphorylation (AAK^T311Ph^) relative to FLAG at centrosomes of metaphase *Drosophila* S2 cells depleted of endogenous AAK and expressing FLAG-AAK^WT^ or FLAG-AAK^K183I^ in the indicated conditions, as presented in (C). All values were normalized to the mean value determined for cells expressing FLAG-AAK^WT^ and endogenous levels of MPS1, which was set to 1. (E) Representative immunofluorescence images of AAK T311 phosphorylation (AAK^T311Ph^) in prometaphase or metaphase *Drosophila* S2 cells depleted of endogenous AAK and expressing EGFP-AAK^WT^ or EGFP-AAK^T360A^ in the indicated conditions. Insets depict magnifications of selected centrosomes. EGFP-AAK^WT^ cells expressing endogenous MPS1 or overexpressing mCherry-MPS1 (OE) were imaged in the same coverslip. EGFP-AAK^T360A^ cells expressing endogenous MPS1 or overexpressing mCherry-MPS1 (OE) were imaged in the same coverslip. (F) Quantification of the levels of AAK T311 phosphorylation (AAK^T311Ph^) relative to EGFP at centrosomes of prometaphase or metaphase *Drosophila* S2 cells depleted of endogenous AAK and expressing EGFP-AAK^WT^ or EGFP-AAK^T360A^ in the indicated conditions, as presented in (E). All values were normalized to the mean value determined for prometaphase cells expressing EGFP-AAK^WT^ and endogenous levels of MPS1, which was set to 1. Data information: Data in (B) are presented as mean ± SD and in (D) and (F) are presented as median ± interquartile range. Statistical analysis in (D) and (F) was performed using a Kruskal– Wallis test for multiple comparisons. P values: ns, not significant; *< 0.05; **< 0.01; ***< 0.001; ****< 0.0001. Scale bar: 5 μm. See also Figure S2.

A series of cell-based assays were next employed to test whether MPS1 stimulates AAK auto-phosphorylation of the T-loop (AAK^T311Ph^) via phosphorylation of T360 in S2 cells. Consistent with the FRET results, over-expressing (OE) MPS1 caused a significant increase in AAK^T311Ph^ at metaphase centrosomes relative to controls (Figures S2C and S2D). Loss of AAK activation upon MPS1 OE in AAK depleted cells (Figures S2C and S2D) could be rescued by expression of WT but not kinase-dead (K183I) AAK (Figures 2C and 2D). Importantly, expression of an AAK non-phosphorylatable T360A mutant (AAK^T360A^), predicted to have a similar structure as AAK^WT^ (Figure S2E), also failed to rescue MPS1-mediated activation of AAK in either prometaphase or metaphase cells (Figures 2E and 2F). Thus, phosphorylation of the conserved AAK T360 residue by MPS1 promotes auto-phosphorylation of the AAK T-loop (T311) and hence efficient kinase activation *in vitro* and in cells.

### Activation of AAK by MPS1 is required for efficient error correction and chromosome congression

The functional contributions of proximity-based activation of AAK by MPS1 to cell division was evaluated by live- and fixed-cell analyses of mitotic S2 cells. To isolate potential chromosome alignment and error correction functions of MPS1 from its role as a checkpoint kinase in *D. melanogaster* [23], cells were arrested in mitosis via RNAi of the APC/C subunit CDC27 [24–26]. Cells efficiently aligned their chromosomes on a metaphase plate upon depletion of CDC27, however, co-depleting MPS1 impaired timely chromosome congression and increased the abundance of persistent polar chromosomes (Figures 3A, 3C and Video S1). The congression defect in CDC27/MPS1 co-depleted cells was rescued by centrosomal targeting (via C-terminal fusions to CM2) of either wild type MPS1 (MPS1^WT^-CM2) or a phospho-mimetic mutant of AAK in which the previously identified MPS1 site (T360) was mutated to aspartic acid (AAK^T360D^-CM2) (Figures 3A, 3C and Video S2). The rescue of chromosome alignment required AAK kinase activity since cells expressing a kinase dead (K183I) version of AAK^T360D^-CM2 (AAK^K183I/T360D^-CM2) failed to effectively congress their chromosomes (Figures 3A, 3C and Video S2). As expected, co-depletion of AAK and CDC27 delayed chromosome congression (Figure 3B, 3D and Video S3). This defect was rescued by the expression of EGFP-AAK^WT^ but not by the non-phosphorylatable T360A AAK mutant (EGFP-AAK^T360A^) (Figure 3B, 3D and Video S3). These results support the conclusion that MPS1-mediated phosphorylation of AAK T360 is required for efficient chromosome congression

**FIGURE 3.**
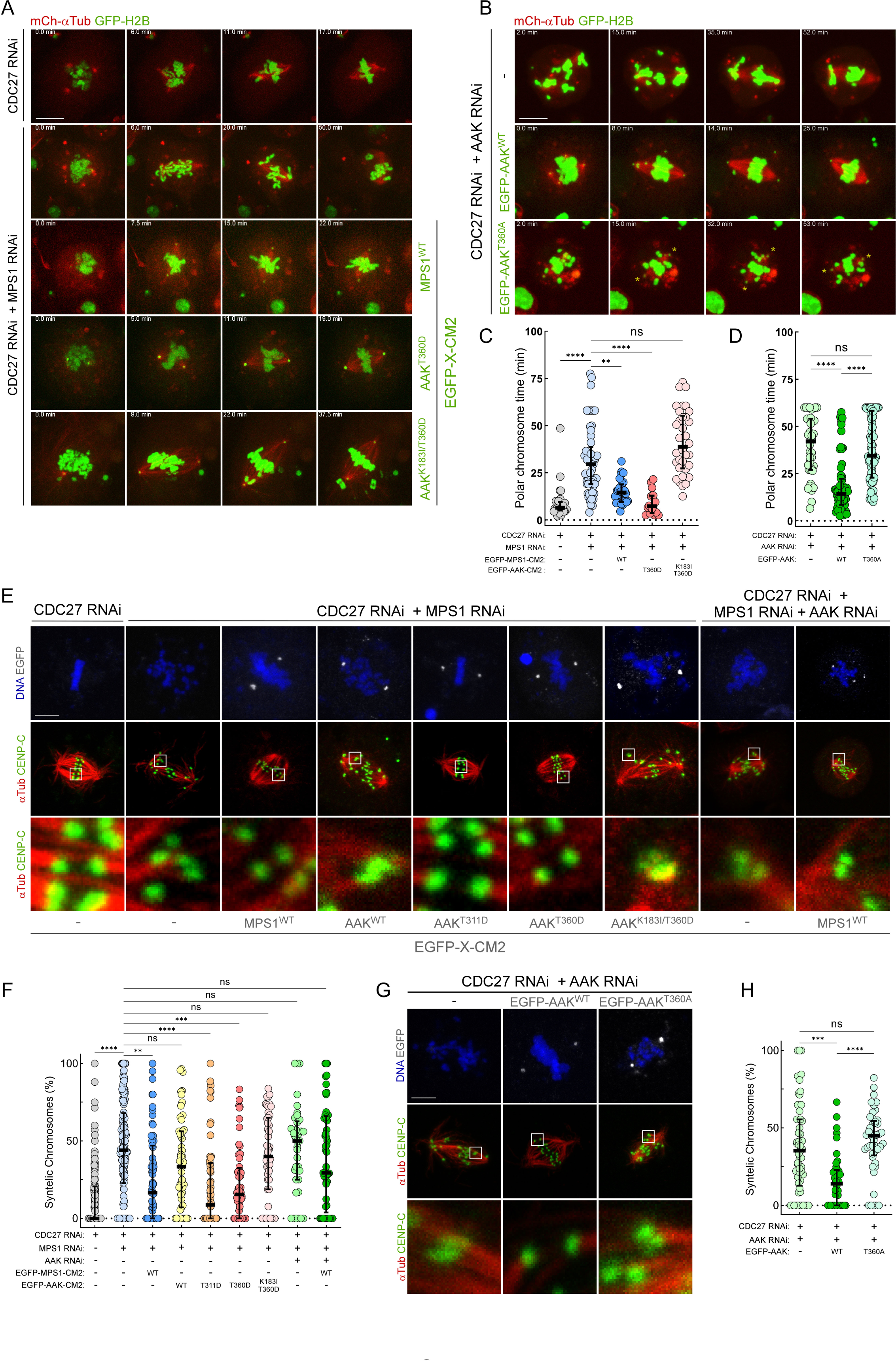
Activation of AAK by MPS1 is required for efficient error correction and chromosome congression. (A) Selected stills of representative mitotic progression movies of CDC27-depleted or CDC27/MPS1 co-depleted *Drosophila* S2 cells stably expressing GFP-H2B, mCherry-αTubulin and the indicated EGFP-tagged and CM2-fused transgenes (EGFP-X-CM2). (B) Selected stills of representative mitotic progression movies of CDC27/AAK co-depleted *Drosophila* S2 cells stably expressing GFP-H2B, mCherry-αTubulin and the indicated transgenes. Asterisks indicates the centrosomes in the EGFP-AAK^T360A^ condition. (C) Quantifications of the congression time of individual polar chromosomes (time each chromosome was retained near the spindle pole) in the conditions indicated in (A). (D) Quantifications of the congression time of individual polar chromosomes (time each chromosome was retained near the spindle pole) in the conditions indicated in (B). (E) Representative immunofluorescence images of mitotic *Drosophila* S2 cells expressing the indicated EGFP-tagged and CM2-fused transgenes (EGFP-X-CM2) and depleted of the indicated proteins. Insets depict magnifications of selected sister-kinetochores in amphitelic or syntelic attachment configurations. (F) Percentage of syntelic kinetochore-microtubule attachments per cell in the indicated conditions, as presented in (E). (G) Representative immunofluorescence images of CDC27/AAK co-depleted *Drosophila* S2 cells expressing EGFP-tagged AAK^WT^ or AAK^T360A^. Insets depict magnifications of selected sister-kinetochores in amphitelic or syntelic attachment configurations. (H) Percentage of syntelic kinetochore-microtubule attachments per cell in the indicated conditions, as presented in (G). Data information: Data in (C), (D), (F) and (H) are presented as median ± interquartile range. Statistical analysis was performed using a Kruskal–Wallis test for multiple comparisons. P values: ns, not significant; *< 0.05; **< 0.01; ***< 0.001; ****< 0.0001. Scale bar: 5 μm. See also Figure S3 and Videos S1, S2 and S3.

The attachment status of kinetochores was next assessed by analyzing confocal Z-sections of chromosomes (DAPI), kinetochores (CENP-C), and microtubules (α-Tubulin) in calcium-treated fixed cells to assess error correction (Figures 3 E-H and S3A-D). Chromosomes were scored as (i) bioriented when each sister kinetochore was clearly associated with microtubules oriented toward opposing spindle poles; (ii) syntelic when sister kinetochore pairs were clearly attached to microtubules from the same spindle pole; and (iii) pinned when the kinetochore pair was within 1 micron of the spindle pole regardless of whether its attachment status could be clearly discerned. While a vast majority of chromosomes were bioriented in CDC27-depleted cells, co-depletion of CDC27 and MPS1 resulted in a significant increase in the frequency of syntelic and pinned chromosomes revealing that MPS1 is required for error correction in fly cells (Figures 3E, 3F, S3A and S3C). The error correction defect in CDC27/MPS1 RNAi cells could be rescued by expressing centrosome-tethered MPS1^WT^-CM2 but not AAK^WT^-CM2 (Figures 3E, 3F, S3A and S3C). Co-depletion of AAK hindered MPS1^WT^-CM2 capacity to promote chromosome alignment and biorientation, further supporting the conclusion that MPS1 acts upstream of AAK in the pole-based error correction pathway (Figures 3E, 3F, S3A and S3C). The requirement for MPS1 to potentiate AAK-mediated error correction could be bypassed in CDC27/MPS1 co-depleted cells by centrosomal targeting of either the T-loop (T311D) or C-lobe (T360D) AAK-CM2 phospho-mimetic mutants but not the catalytic dead AAK^K183I/T360D^-CM2 version (Figures 3E, 3F, S3A and S3C).

To directly test the requirement of AAK T360 phosphorylation for error correction, we co-depleted CDC27 and endogenous AAK and evaluated kinetochore-microtubule attachments upon the expression of untethered versions of AAK^WT^ and AAK^T360A^. Contrasting with EGFP-AAK^WT^, the non-phosphorylatable EGFP-AAK^T360A^ mutant failed to restore chromosome biorientation and alignment in S2 cells depleted of endogenous AAK (Figures 3G, 3H, S3B and S3D). These findings support the conclusion that pole-based error correction in fly cells is carried out by AAK activated via phosphorylation of its C-lobe (T360) by MPS1.

### Proximity-based activation of AAK by MPS1 is conserved in human cells

To examine if proximity-based activation of AAK by MPS1 is conserved outside *D. melanogaster,* the extent of AAK activation was evaluated under experimental conditions where the positioning of kinetochores relative to centrosomes/spindle poles could be manipulated in human RPE-1 cells [27]. When compared to control monopolar spindles (STLC treatment), kinetochores were positioned closer to the monopole center upon inhibition of the kinetochore-associated motor CENP-E with GSK923295 (CENP-Ei) while depletion of the Dynein adaptor Spindly (siSpindly) positioned kinetochores further from the monopole center (Figures 4A-C). Consistent with there being spatial cross-talk between kinetochores and centrosomes, AAK activation (AAK^T288Ph^) levels at centrosomes/spindle poles was significantly increased in the CENP-Ei cells and significantly reduced in the siSpindly cells relative to control cells (Figures 4D and 4E). Active AAK^T288Ph^ signal at centrosomes was significantly reduced compared to controls in nocodazole-(Figures S4A and S4B) and STLC-treated cells (Figure 4F and 4G) when MPS1 was inhibited with Cpd-5 (MPS1i). Importantly, inhibiting MPS1 prevented the increase in AAK activation in CENP-Ei monopoles (Figure 4F and 4G). Thus, the potentiation in AAK activation observed when kinetochores were positioned closer to centrosomes in human RPE-1 cells required MPS1 activity.

**Figure 4.**
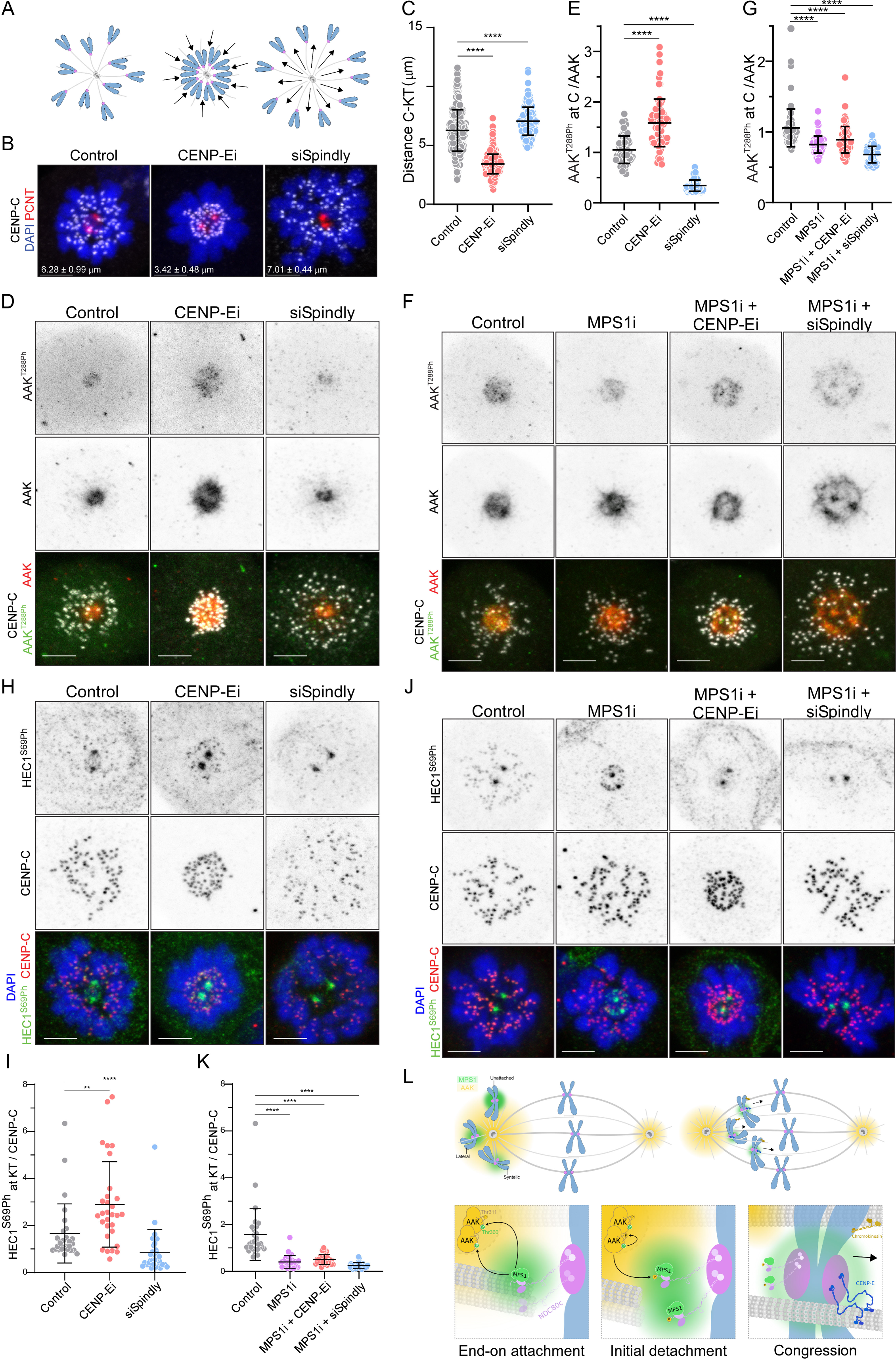
Proximity-based activation of AAK by MPS1 is conserved in human cells. (A) Schematic representation of the monopolar spindle assay used to assess AAK^T288Ph^ and HEC1^S69Ph^ changes in response to the distance between centrosomes and kinetochores. (B) Representative immunofluorescence images of STLC-treated RPE-1 cells in control, CENP-E inhibited (CENP-Ei) and Spindly depleted (siSpindly) conditions. (C) Quantification of centrosome (PCNT)-kinetochore (CENP-C) (C-KT) distances of STLC-treated RPE-1 cells in the indicated conditions, as presented in (B). (D) Representative immunofluorescence images of AAK T288 phosphorylation (AAK^T288Ph^) and AAK levels in STLC-treated RPE-1 cells in control, CENP-E inhibited (CENP-Ei) and Spindly depleted (siSpindly) conditions. (E) Quantification of the levels of AAK T288 phosphorylation (AAK^T288Ph^) relative to total AAK levels at centrosomes of STLC-treated RPE-1 cells in the indicated conditions, as presented in (D). All values were normalized to the median value determined for control cells, which was set to 1. (F) Representative immunofluorescence images of AAK T288 phosphorylation (AAK^T288Ph^) and AAK levels in STLC-treated RPE-1 cells in control, MPS1 inhibited (MPS1i), MPS1/CENP-E co-inhibited (MPS1i+CENP-Ei) and MPS1 inhibited in Spindly depleted background (MPS1i+siSpindly). In all conditions the proteasome was inhibited with MG132. (G) Quantification of the levels of AAK T288 phosphorylation (AAK^T288Ph^) relative to total AAK levels at centrosomes of STLC-treated RPE-1 cells in the indicated conditions, as presented in (F). All values were normalized to the median value determined for control cells, which was set to 1. (H) Representative immunofluorescence images of HEC1 S69 phosphorylation (HEC1^S69Ph^) and CENP-C levels in STLC-treated RPE-1 cells in control, CENP-E inhibited (CENP-Ei) and Spindly depleted (siSpindly) conditions. (I) Quantification of the levels of HEC1 S69 phosphorylation (HEC1^S69Ph^) relative to CENP-C levels at kinetochores of STLC-treated RPE-1 cells in the indicated conditions, as presented in (H). All values were normalized to the median value determined for control cells, which was set to 1. (J) Representative immunofluorescence images of HEC1 S69 phosphorylation (HEC1^S69Ph^) and CENP-C levels in STLC-treated RPE-1 cells in control, MPS1 inhibited (MPS1i), MPS1/CENP-E co-inhibited (MPS1i+CENP-Ei) and MPS1 inhibited in Spindly depleted background (MPS1i+siSpindly). In all conditions the proteasome was inhibited with MG132. (K) Quantification of the levels of HEC1 S69 phosphorylation (HEC1^S69Ph^) relative to CENP-C levels at kinetochores of STLC-treated RPE-1 cells the indicated conditions, as presented in (J). All values were normalized to the median value determined for control cells, which was set to 1. (L) Proposed model for proximity-based activation of AAK by MPS1 as a strategy to potentiate error correction of polar chromosomes and ensure chromosome congression. A pool of MPS1 recruited to kinetochores of polar chromosomes indirectly contributes to pole-based error correction by phosphorylating AAK on the conserved residue T360. This potentiates AAK activation at centrosomes and thereby its capacity to phosphorylate the N-terminal tail of HEC1/NDC80 at kinetochores to destabilize end-on attachments. This mechanism corrects and prevents syntelic kinetochore-microtubule interactions in the vicinity of centrosomes and ensures efficient chromosome congression. Data information: Data are presented in (C), (E), (G), (I) and (K) as mean ± SD. Statistical analysis was performed using a Mann-Whitney non-parametric t-test. P values: ns, not significant; *< 0.05; **< 0.01; ***< 0.001; ****< 0.0001. Scale bar 5 mm. N ≥ 75 cells for each condition. See also Figure S4.

AAK targets a number of the same sites on the N-terminal tail of NDC80 as ABK but S69 on HEC1 (human NDC80) is preferentially phosphorylated by AAK [28]. Thus, the extent of S69 phosphorylation (HEC1^S69Ph^) was measured as a function of kinetochore positioning within monopolar spindles. Consistent with the extent of AAK activation (AAK^T288Ph^ levels), phosphorylation of HEC1-S69 was increased in CENP-Ei cells and reduced in siSpindly cells relative to controls (Figures 4H and 4I). In nocodazole-treated cells, HEC1^S69Ph^ levels were reduced to the same extent following inhibition of either MPS1 or AAK and to a lesser extent upon ABK inhibition reflecting an upstream role of MPS1 in the preferential phosphorylation of HEC1-S69 by AAK (Figures S4C and S4D). Importantly, HEC1^S69Ph^ levels were significantly reduced when MPS1 was inhibited including in CENP-Ei cells that normally exhibited a significant increase in HEC1^S69Ph^ levels (Figures 4J and 4K). Altogether, the data support the conclusion that kinetochore proximity-based activation of AAK by MPS1 potentiates phosphorylation of the bona-fide error correction substrate HEC1 in human cells.

## DISCUSSION

We propose a model whereby a proximity-dependent crosstalk between kinetochores and centrosomes/spindle poles promotes error correction (Figure 4L). The model envisions that as erroneous kinetochore-microtubule interactions (e.g. a syntelic attachment) become positioned within the vicinity (∼1 μm) of the spindle pole, kinetochore-recruited MPS1 phosphorylates AAK on a conserved site (T360 in flies) located in the αG-helix of the kinase C-lobe. Phosphorylation of the C-lobe, in turn, triggers auto-catalytic phosphorylation of the AAK T-loop to amplify a ∼2 μm radius activity gradient [10,11,29] to locally promote microtubule detachment via phosphorylation of kinetochore substrates such as the NDC80 complex. AlphaFold3 predicts that phosphorylation of T360 causes an outward stretch of the αG-helix that pulls on the T-loop by interacting with D317 in manner that could render the T-loop more flexible and leave the activating T311 more exposed for phosphorylation (Figure S2F-H). Further investigation into the structural basis of this novel activation mechanism is warranted.

It has been proposed that the establishment of stable end-on attachments prevents recruitment of MPS1 to kinetochores because of mutually exclusive binding to either microtubules or MPS1 by the CH-domains of the NDC80 complex [13–15]. However, a pool of MPS1 still localizes to microtubule-attached kinetochores either through association with the NDC80 complex or other kinetochore receptors [7,12]. Three recent studies in budding yeast showed that MPS1 binds to regions in the CH-domains of the NDC80 complex that are on the opposite side of the microtubule binding region [16–18] and, accordingly, purified budding yeast NDC80 complex was shown to simultaneously bind microtubules and MPS1 *in vi*tro [17]. It has been suggested that this interaction interface is conserved [18]; if so then this mechanism could explain how a population of MPS1 remains at microtubule-attached metazoan kinetochores. Regardless of how it is recruited, we hypothesize that the residual MPS1 population on erroneously attached polar kinetochores activates AAK to promote their own correction.

An interesting feature of the crosstalk pathway described here is that it contains an intrinsic positive feedback loop since the destabilization of kinetochore-microtubule interactions by AAK-mediated phosphorylation of NDC80 would recruit more MPS1. Such a spatio-chemical feedback loop would heavily disfavor production of end-on attached kinetochores in the vicinity of spindle poles. Thus, we speculate that MPS1-driven potentiation of AAK activity following nuclear envelope breakdown is important to minimize the formation of end-on attachments near the centrosomes, where syntelic attachments would otherwise be prone to accumulate due to the high density of nucleated microtubules. Instead, robust activation of AAK favors the formation of lateral attachments and facilitates motor activation of CENP-E [30] to ensure timely chromosome congression towards the cell equator, where amphitelic attachments can form synchronously as a result of interactions between non-centrosomal short microtubules emanating from kinetochores and bundles of antiparallel microtubules [31–33].

Overall, our findings support the conclusion that a pool of MPS1 at kinetochores indirectly contributes to pole-based error correction. This model is broadly consistent with an earlier proposal that MPS1 plays an indirect role in error correction with a notable difference being that the relevant downstream target of MPS1 is AAK rather than ABK [1–3]. MPS1 clearly activated AAK in our cell-based assays, but consistent with prior findings [4–6] we did not see evidence of MPS1-mediated ABK activation in cells despite the facts that (i) the centromere/kinetochore localized ABK is closer than centrosomal AAK to kinetochore-bound MPS1, (ii) the C-lobe threonine is present in ABK αG-helix, and (iii) ABK is phosphorylated and activated by MPS1 *in vitro* (not shown). Since constitutive ABK activation would be detrimental to forming stable attachments, kinetochore/centromere associated protein phosphatases likely protect ABK from MPS1-mediated activation in cells. Direct and indirect roles for MPS1 in error correction need not be mutually exclusive; in fact, it now appears that diverse organisms rely on each mechanism to different extents in carrying out error correction. Budding yeast utilize MPS1 to directly target the NDC80 complex without indirectly inducing error correction through activation of AURORA kinase [9]. Conversely, our work shows that activation of AAK-mediated error correction by kinetochore-recruited MPS1 is the dominant pathway in flies although a direct role for MPS1 in error correction cannot be ruled out. Since we observed proximity-based activation of AAK by MPS1 in RPE-1 cells, efficient error correction in humans likely requires both indirect and direct pathways via MPS1-mediated phosphorylation of AAK and kinetochore-microtubule attachment factors respectively. Future efforts to dissect the relative contributions of direct versus indirect roles of MPS1 in error correction are warranted, but the findings presented here define a novel and conserved mechanism by which polar kinetochores modulate centrosome-based activities to promote accurate chromosome segregation.

## ACKNOWLEDGMENTS

We thank Geert Kops, Jennifer DeLuca, Mónica Bettencourt-Dias, René Medema, Christian Lehner and Scott Hawley for reagents. We thank Andrea Musacchio for critically discussing the data. Work in C.C. and C.E.S. lab was funded by National Funds through FCT—Fundação para a Ciência e a Tecnologia, I.P., under the project UIDB/04293/2020 and by the project “Cancer Research on Therapy Resistance: From Basic Mechanisms to Novel Targets” - NORTE-01-0145-FEDER-000051, supported by Norte Portugal Regional Operational Programme (NORTE 2020), under the PORTUGAL 2020 Partnership Agreement, through the European Regional Development Fund (ERDF). C.C. is supported by a Scientific Employment Stimulus contract (2020.00067.CEECIND) from Fundação para a Ciência e a Tecnologia. N.L. (SFRH/BD/136526/2018) and M.M. (SFRH/BD/123306/2016) were supported by PhD fellowships from Fundação para a Ciência e a Tecnologia. This work was supported by an NIH grant (GM107026) to T.J.M. International collaboration was supported with Fulbright Awards to T.J.M and N.L. Work in T.B. lab was supported by funding from the Deutsche Forschungsgemeinschaft (DFG) (project number: 5041140321). Work in M.B. lab was supported by the Novo Nordisk Foundation (NNF19OC0058504), the Independent Research Fund Denmark (3101-00075B), and the Lundbeck Foundation (R434-2023-431). The authors acknowledge the i3S Scientific Platform ALM, member of the national infrastructure Portuguese Platform of Bioimaging for excellent support.

## AUTHOR CONTRIBUTIONS

C.C. conceptualized the study and T.J.M. wrote the original manuscript draft with contributions from C.C.; C.C. and T.J.M. coordinated the work; C.C. designed the experiments, with contributions from N.L., C.E.S., M.B. and T.J.M.; Data acquisition: N.L. performed the major experiments presented in Figures 1, 2, 3, S1, S2 and S3, with contributions from F.B., M.M., and I.P.; S.R.C. performed the experiments presented in Figures 4 and S4; T.B. performed the mass spectrometry analysis and P.D.P. generated the Alphafold models. Data analysis and interpretation: N.L., F.B., M.M., P.D.P., I.P., A.V., T.B., C.E.S., M.B., T.J.M. and C.C.; Supervision: C.E.S., M.B., T.J.M. and C.C. Funding acquisition C.E.S., T.B., M.B., T.J.M. and C.C.; N.L. assembled Figures 1, 2, 3, S1, S2, S3 and Videos S1, S2 and S3; S.R.C. and M.B. assembled Figures 4 and S4. All authors reviewed the manuscript and agreed with its contents.

## SUPPLEMENTAL INFORMATION

**Document S1.** Figures S1–S4 and Video legends

**Video S1.** Depletion of MPS1 delays chromosome congression, related to Figures 3A and 3C

**Video S2.** Centrosomal-tethered MPS1 or a AAK^T360D^ mutant can restore chromosome congression in MPS1-depleted cells, related to Figure 3A and 3C

**Video S3.** Phosphorylation of AAK T360 is required for chromosome congression, related to Figure 3B and 3D

## SUPPLEMENTAL FIGURE LEGENDS

**FIGURE S1.**
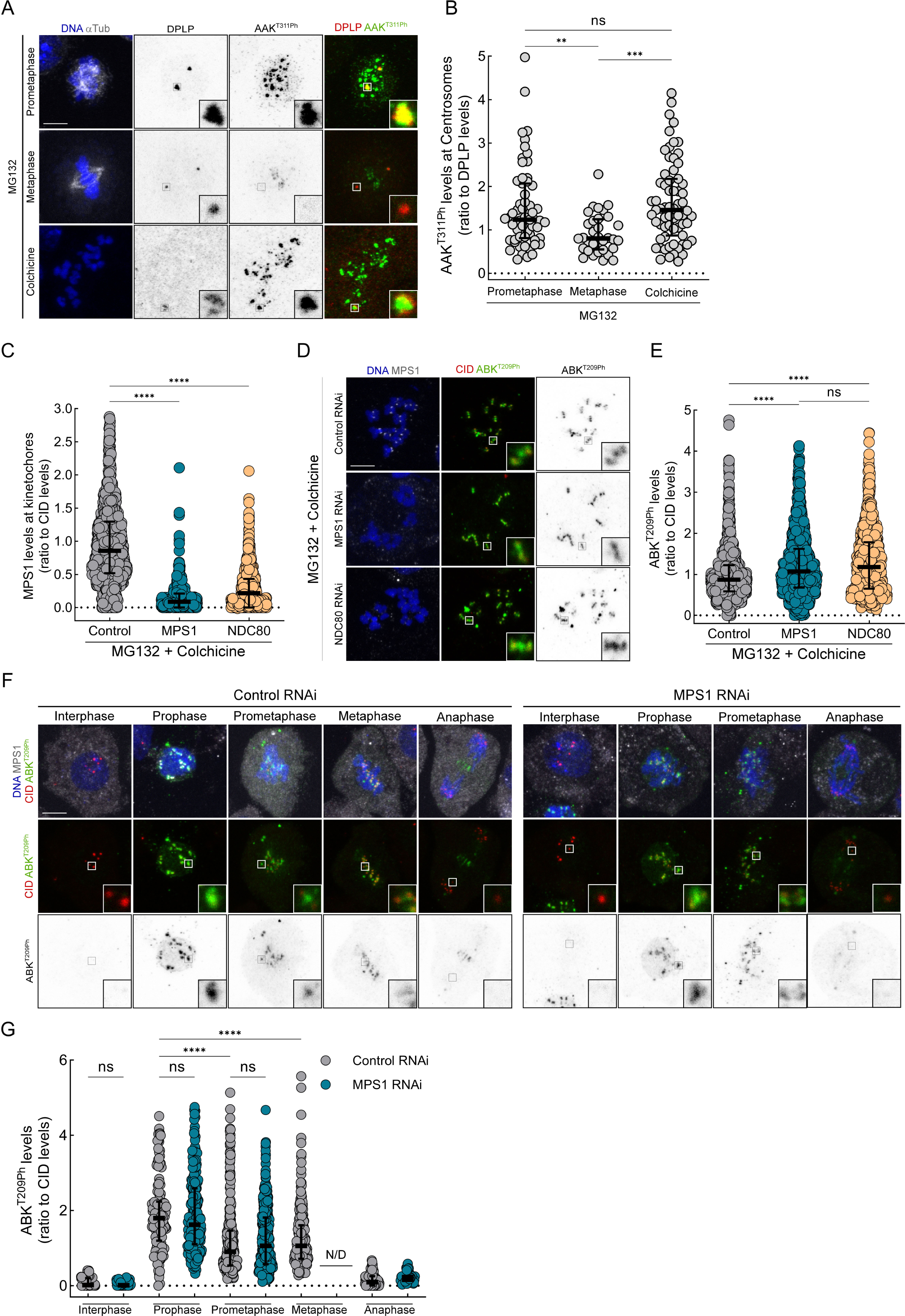
MPS1 is required for mitotic activation of AAK at centrosomes but dispensable for ABK activation at centromeres. Related to Figure 1. (A) Representative immunofluorescence images of AAK T311 phosphorylation (AAK^T311Ph^) levels in MG132-treated *Drosophila* S2 cells in the indicated conditions. Insets depict magnifications of selected centrosomes. (B) Quantification of the levels of AAK T311 phosphorylation (AAK^T311Ph^) relative to DPLP at centrosomes in the indicated conditions, as presented in (A). All values were normalized to the mean value determined for prometaphase control cells treated with MG132, which was set to 1. (C) Quantification of the levels of total MPS1 levels relative to CID at kinetochores in MG132 and colchicine-treated *Drosophila* S2 cells in the indicated conditions, as presented in (D). All values were normalized to the mean value determined for control cells treated with MG132 and colchicine, which was set to 1. (D) Representative immunofluorescence images of ABK T209 phosphorylation (ABK^T209Ph^) and MPS1 levels in MG132 and colchicine-treated *Drosophila* S2 cells in the indicated conditions. Insets depict magnifications of selected kinetochores. (E) Quantification of the levels of ABK T209 phosphorylation (ABK^T209Ph^) relative to CID at kinetochores/centromeres in the indicated conditions, as presented in (D). All values were normalized to the mean value determined for control cells treated with MG132 and colchicine, which was set to 1. (F) Representative immunofluorescence images of ABK T209 phosphorylation (ABK^T209Ph^) in control and MPS1-depleted *Drosophila* S2 cells in the indicated cell cycle stages. Insets depict magnifications of selected kinetochores/centomeres. (G) Quantification of the levels of ABK T209 phosphorylation (ABK^T209Ph^) relative to CID at kinetochores/centromeres of control and MPS1-depleted *Drosophila* S2 cells in the indicated cell cycle stages, as presented in (F). All values were normalized to the mean value determined for prometaphase control cells, which was set to 1. Data information: Data in (B), (C), (E) and (G) are presented as median ± interquartile range. Statistical analysis was performed using a Kruskal–Wallis test for multiple comparisons. P values: ns, not significant; *< 0.05; **< 0.01; ***< 0.001; ****< 0.0001. Scale bar: 5 μm.

**FIGURE S2.**
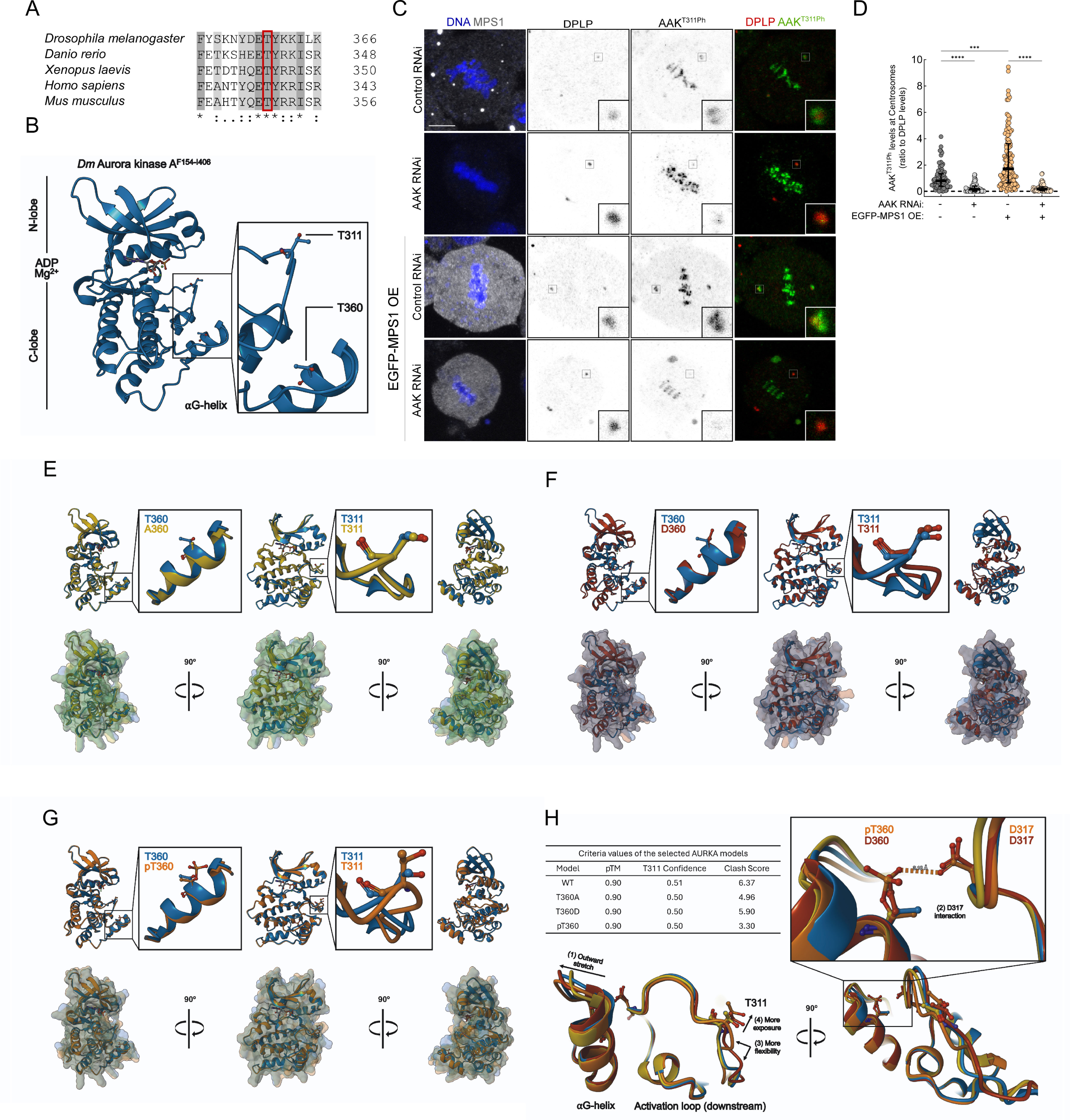
Phosphorylation of AAK T360 by MPS1 potentiates AAK T-loop autophosphorylation. Related to Figure 2. (A) Clustal Omega (EMBL-EBI) local sequence alignment for the indicated AAK orthologues. Residues conservation: (*), fully conserved; (:), strongly similar properties; (.), weakly similar properties. MPS1-mediated phosphorylation of Threonine 360 (T360) in *Drosophila* AAK was identified by MS analysis after *in vitro kinase* assays. The identified residue is conserved in AAK orthologues (red rectangular box). (B) AlphaFold 3 model of the kinase domain (154-406aa) of *Drosophila melanogaster* AAK (Q9VGF9) in crystal-like conditions (ADP and two Mg2+ ions). (C) Representative immunofluorescence images of AAK T311 phosphorylation (AAK^T311Ph^) levels in metaphase *Drosophila* S2 cells in the indicated conditions. Insets depict magnifications of selected centrosomes. (D) Quantification of the levels of AAK T311 phosphorylation (AAK^T311Ph^) relative to DPLP at centrosomes of metaphase *Drosophila* S2 cells in the indicated conditions, as presented in (C). All values were normalized to the mean value determined for control cells expressing endogenous levels of MPS1, which was set to 1. (E) Superposed AlphaFold 3 models of *Drosophila* AAK^WT^ (cyan) and AAK^T360A^ (yellow) showing that the T360A mutation is not predicted to induce structural changes on the ⍺G-helix, nor on the T-loop. (F) Superposed AlphaFold 3 models of *Drosophila* AAK^WT^ (cyan) and AAK^T360D^ (red) showing that the T360D mutation is predicted to cause a displacement in the ⍺G-helix, opening room for more flexibility of the T-loop and exposure of T311. (G) Superposed AlphaFold 3 models of *Drosophila* AAK^WT^ (cyan) and AAK^WT^ phosphorylated in T360 (AAK^pT360^) (orange) predicting that the phospho-threonine causes steric hindrance leading to ⍺G-helix displacement and increased T-loop flexibility and exposure. (H) The table shows the overall structure pTM, T-loop Confidence and ClashScore for the best models and underneath is presented the predicted model by which phosphorylation of T360 facilitates AAK T311 autophosphorylation. This model consists of four steps, (1) an outward stretch of the ⍺G-helix that pulls on the T-loop by (2) interacting with D317 in manner that could (3) render the T-loop more flexible and (4) leave the activating T311 more exposed for phosphorylation. Data information: Data in (D) are presented as median ± interquartile range. Statistical analysis was performed using a Kruskal–Wallis test for multiple comparisons. P values: ns, not significant; *< 0.05; **< 0.01; ***< 0.001; ****< 0.0001. Scale bar: 5 μm.

**FIGURE S3.**
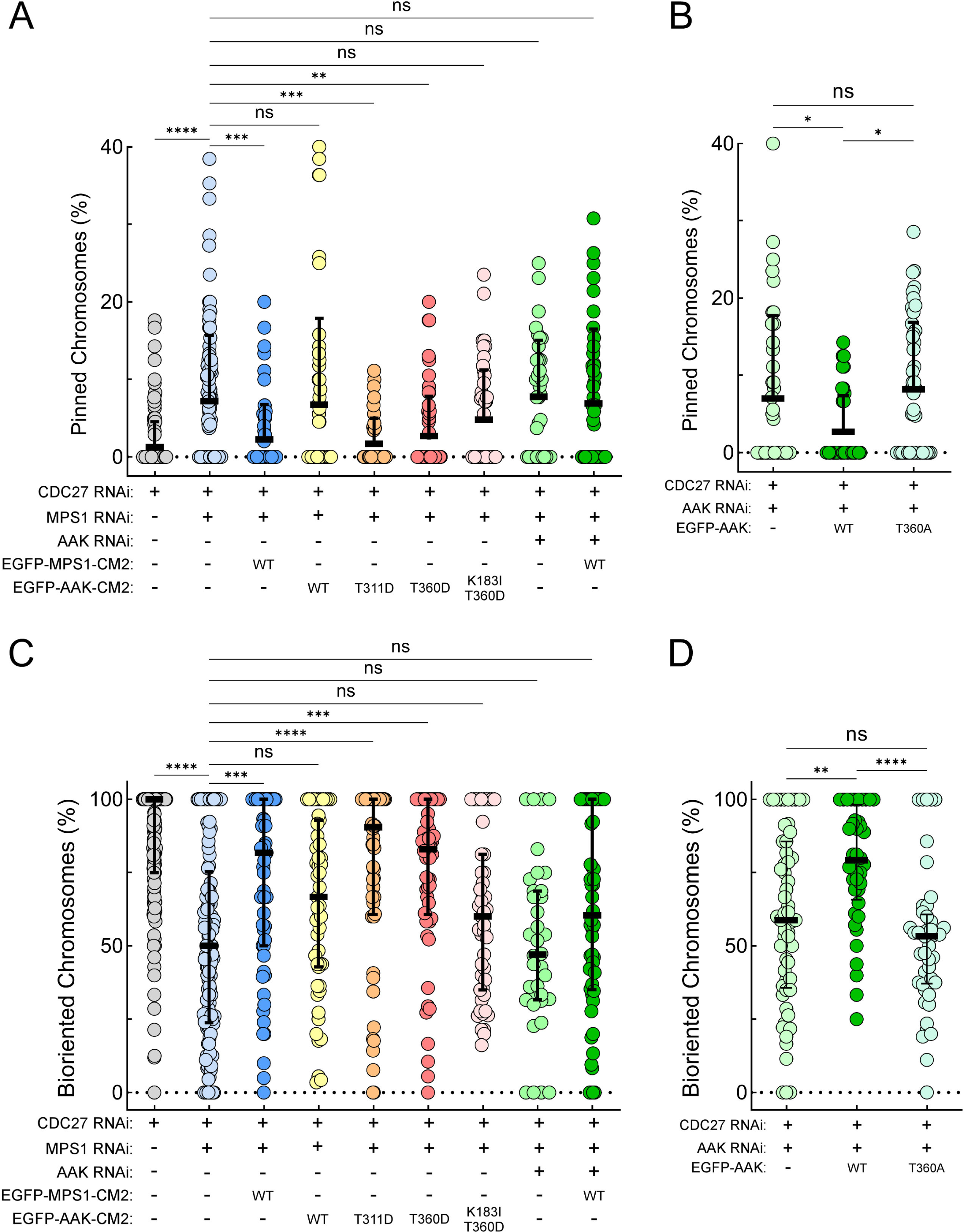
Activation of AAK by MPS1 is required for chromosome congression and biorientation in *Drosophila* S2 cells. Related to Figure 3. (A) Percentage of chromosomes pinned to centrosomes/spindle poles per *Drosophila* S2 cell in the indicated conditions, as presented in Figure 3E. (B) Percentage of chromosomes pinned to centrosomes/spindle poles per *Drosophila* S2 cell in the indicated conditions, as presented in Figure 3G. (C) Percentage of bioriented chromosomes per *Drosophila* S2 cell in the indicated conditions, as presented in Figure 3E. (D) Percentage of bioriented chromosomes per *Drosophila* S2 cell in the indicated conditions, as presented in Figure 3G. Data information: Data in (A) and (B) are presented as mean ± SD; and in (C) and (D) as median ± interquartile range. Statistical analysis was performed using a Kruskal–Wallis test for multiple comparisons. P values: ns, not significant; *< 0.05; **< 0.01; ***< 0.001; ****< 0.0001.

**Figure S4.**
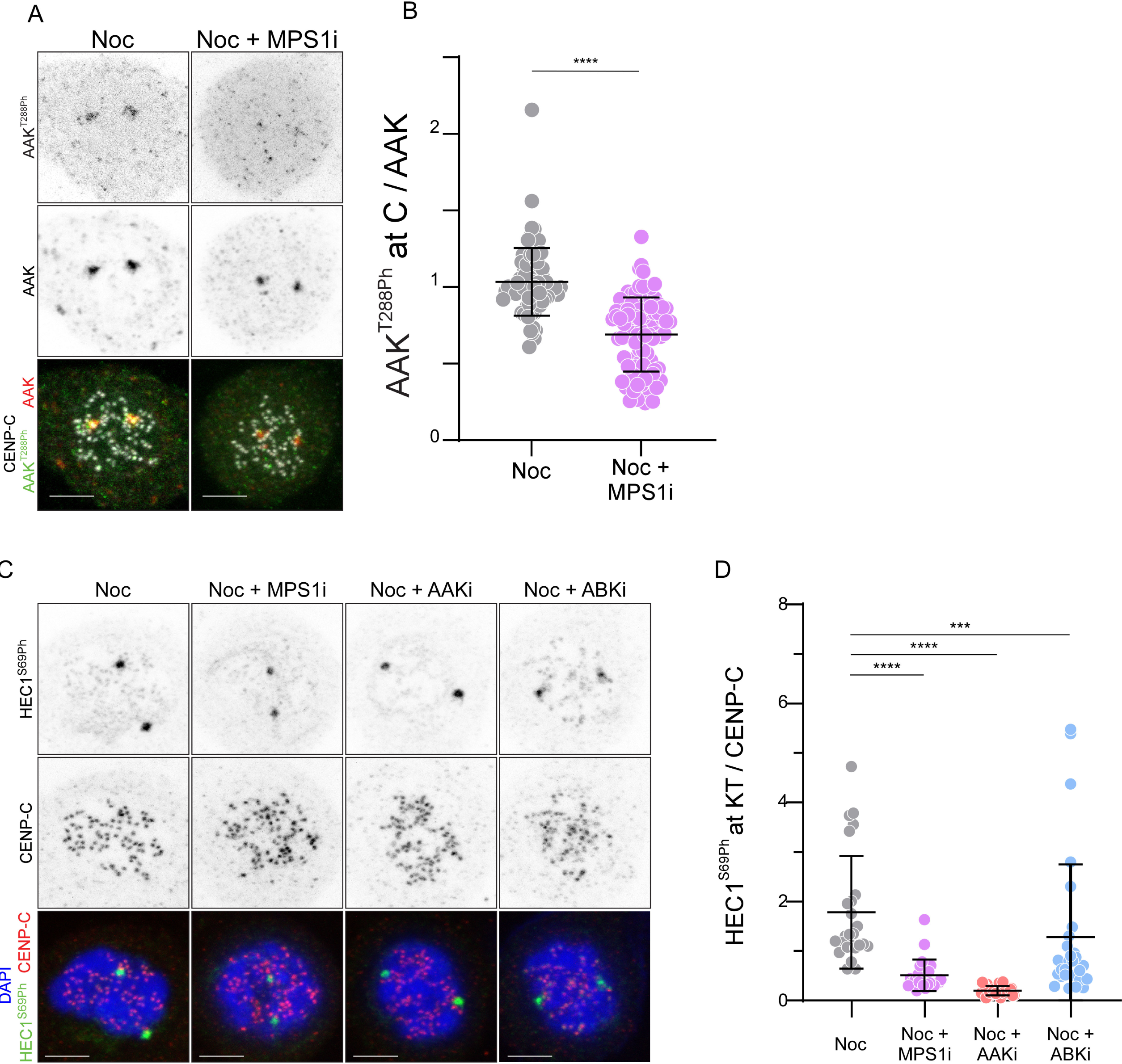
Inhibition of MPS1 impairs AAK activation at centrosomes and HEC1-69 phosphorylation at unattached kinetochores in RPE-1 cells. Related to Figure 4. (A) Representative immunofluorescence images of AAK T288 phosphorylation (AAK^T288Ph^) and AAK levels in nocodazole-treated RPE-1 cells in control and MPS1-inhibited (MPS1i) conditions. Proteasome was inhibited by MG132 to prevent mitotic exit after MPS1 inhibition. (B) Quantification of the levels of AAK T288 phosphorylation (AAK^T288Ph^) relative to total AAK levels at centrosomes of nocodazole-treated RPE-1 cells in the indicated conditions as presented in (A). All values were normalized to the median value determined for control cells, which was set to 1. (C) Representative immunofluorescence images of HEC1 S69 phosphorylation (HEC1^S69Ph^) and CENP-C levels in nocodazole-treated RPE-1 cells in control, MPS1-inhibited (MPS1i), AAK-inhibited (AAKi) and ABK-inhibited (ABKi) conditions. Proteasome was inhibited by MG132 to prevent mitotic exit after MPS1 inhibition. (D) Quantification of the levels of HEC1 S69 phosphorylation (HEC1^S69Ph^) relative to CENP-C levels at kinetochores of nocodazole-treated RPE-1 cells in the indicated conditions as presented in (C). All values were normalized to the median value determined for control cells, which was set to 1. Data information: Data are presented in (B) and (D) as mean ± SD. Statistical analysis was performed using a Mann-Whitney non-parametric t-test. P values: ns, not significant; *< 0.05; **< 0.01; ***< 0.001; ****< 0.0001. Scale bar 5 mm. N ≥ 272 kinetochores from at least 10 cells for each condition. Individual dots represent kinetochore quantification average for each cell.

## MATERIALS AND METHODS

### *Drosophila* S2 cell cultures, RNAi-mediated depletion and drug treatments

*Drosophila* S2 cells (S2-DGRC) were cultured at 25°C in Schneider’s Insect Medium (Sigma-Aldrich) supplemented with 10% FBS, and dsRNA incubation was performed as previously described [23]. The PCR product used as template to produce MPS1 RNAi was amplified with the set of primers MPS1F (5′-TAATACGACTCACTATAGGGTCTTCCAAACACCTATGACG-3′) and MPS1R (5′-TAATACGACTCACTATAGGGCGTTTAGATATCCCTGCACCA-3′). The PCR product used as template to produce AAK 5′UTR RNAi was amplified with AAK5′_F (5′-TAATACGACTCACTATAGGGAGAACTTGCCATTCGCCTCATC-3′) and AAK5′_R (5′-TAATACGACTCACTATAGGGAGAGACGGCACGGCACAGGCACTCG-3′). The PCR product used as template to produce AAK 3′UTR RNAi was amplified with AAK3′_F (5′-TAATACGACTCACTATAGGGAGAACACATTCTTGTTTAATTTTC-3′) and AAK3′_R (5′-TAATACGACTCACTATAGGGAGAGAAAACACACACAAACTTT-3′). The PCR product used as template to produce NDC80 RNAi was amplified with NDC80F (5′-TAATACGACTCACTATAGGGTTTGATCCGGATGTTTTGTTGTAGTTC-3′) and NDC80R (5′-TAATACGACTCACTATAGGGTTTTTTAATTTGAAAACAAAGAATACTCTT-3′). The PCR product used as template to produce CDC27 RNAi was amplified with CDC27F (5′-TAATACGACTCACTATAGGGAACAATAGC-3′) and CDC27R (5′-TAATACGACTCACTATAGGGTCTTCATGTAGAATTGCATGGC-3′). At selected time points, cells were collected and processed for immunofluorescence or time-lapse microscopy. When required, cells were subjected to several drug treatments before being collected and processed for analysis. To promote microtubule depolymerization, cells were incubated for 2 h with 30 µM colchicine (Sigma-Aldrich). To inhibit the proteasome, cells were incubated for 3 h with 20 µM MG132 (Calbiochem).

### Human RPE1 cell cultures, siRNA-mediated depletion and drug treatments

The immortalized human retinal epithelial cell line hTERT RPE-1 (ATCC) was grown in DMEM/F12 supplemented with 10% FBS and cultured at 37°C in humidified conditions with 5% CO_2_. For siRNA-mediated depletion experiments, cells were transfected in OptiMEM (Thermo Fisher) with Lipofectamine RNAiMAX (Thermo Fisher) with 40 nM siRNAs for 48 h – siSpindly: 5′-GAAAGGGUCUCAAACUGAA-3′, siControl (non-targeting control siRNA): 5′-UGGUUUACAUGUCGACUAA-3′. Where indicated, monopolar cells were induced by inhibiting Eg5 using 5 µM S-Trityl-L-Cysteine (STLC; Santa Cruz Biotechnology) which was added to the medium 5 h before fixation. CENP-E was inhibited by adding 200 nM GSK923295 (MedChemExpress) to the medium 1 h before fixation. MPS1 was inhibited by adding 500 nM Cpd-5 ([22]; a gift from René Medema, Oncode Institute, The Netherlands Cancer Institute, The Netherlands) to the medium 1 h before fixation. To avoid mitotic exit after MPS1 inhibition, the proteasome was inhibited by adding 5 µM MG132 (Sigma) to the medium 1 h before MPS1 inhibition. To depolymerize microtubules, 2 µM nocodazole (Sigma) was added 2 h before fixation. For inhibition of AAK and ABK, 250 nM MLN-8054 (Selleck Chemicals) and 4 µM ZM447439 (AstraZeneca) were respectively added to the cells 2 h before fixation.

### Transgene constructs and S2 cell transfection

EGFP-MPS1^WT^, H2B-GFP and mCherry-α-Tubulin constructs were previously characterized [23]. For the generation of the expression vector mCherry-MPS1^WT^, mCherry cDNA was amplified by PCR and inserted into the EGFP-MPS1 construct, by FastCloning [34], in replacement of previously removed EGFP. EGFP-AAK^WT^ and FLAG-AAK^WT^ constructs were generated by PCR amplification of AAK cDNA and insertion into modified versions of pENTR entry vector through FastCloning [34]. Entry clones were then used for Gateway recombination (Thermo Fischer Scientific) together with pHGW (for EGFP-AAK constructs) or pHFMW (for FLAG-AAK constructs) destination vectors (heat-shock inducible promoter, N-terminal EGFP or FLAG tag, respectively). To generate FLAG-AAK^K183I^, EGFP-AAK^T311D^, EGFP-AAK^T360A^, EGFP-AAK^T360D^ and EGFP-AAK^K183I/T360D^ versions, either FLAG-AAK^WT^ or EGFP-AAK^WT^ expression vectors had their codons corresponding to K183, T311, T360, T360 and K183 and T360 changed by site-directed mutagenesis with a primer harboring the desired mutation to Isoleucine (I), Aspartate (D), Alanine (A), Aspartate (D) and Isoleucine (I) and Aspartate (D), respectively. The PCR reactions were performed with Phusion polymerase (Thermo Fisher Scientific). To generate the EGFP-MPS1^WT^-CM2 and EGFP-AAK-CM2 constructs, the CM2 domain of Centrosomin (CNN) cDNA was amplified by PCR and was inserted in the C-terminal region of the available EGFP-MPS1^WT^ and EGFP-AAK constructs, respectively, by FastCloning [34]. Stable transfection of indicated vectors into S2 cells was performed using Effectene Transfection Reagent (Qiagen, Hilden, Germany), according to the manufacturer’s instructions. To induce EGFP-MPS1^WT^, mCh-MPS1^WT^ and EGFP-MPS1^WT^-CM2 expression, S2 cells were incubated overnight with 200 μM CuSO_4_ at 25°C. To induce the expression of either AAK constructs cells were heat-shocked for 30 min at 37°C and allowed to rest for at least 6 h before being processed for immunofluorescence or live cell imaging.

### FRET-based AURORA kinase sensor

S2 cells stably expressing the CM2-FRET reporter, GFP-H2B and either mCherry-α-tubulin or mCherry-MPS1 were seeded onto Concanavalin A-(ConA) (Sigma-Aldrich) treated 35 mm glass bottom petri dishes (Cellvis) for 1 hour prior to bringing the volume of Schneider’s media up to 2 mL. Cells were then imaged at 24°C on a TiE inverted microscope (Nikon) equipped with an iXON EMCCD camera (Andor Technology) using a 100x 1.4 NA Plan Apo violet-corrected series differential interference contrast objective (Nikon). Metamorph software (Molecular Devices) was used to control the imaging system. Mitotic cells and the best focal plane were identified on the RFP channel (mCherry-α-tubulin or mCherry-MPS1). After acquiring the RFP image, sequential images of mTurquoise2, mVenus, and FRET (ECFP excitation, EYFP emission) were taken with equal exposure times (200 ms). Background-corrected fluorescence intensities for mTurquoise2 and FRET were measured in Metamorph software by subtraction of local background using a duplicate area. The reported FRET emission ratios represent the ratios of the background corrected FRET signal over the background corrected mTurquoise2 signal and values were normalized against the mean prometaphase FRET emission ratio for each biological replicate. Over-expression of MPS1 was induced by overnight incubation with 200 μM CuSO_4_. AAK was inhibited by treating the seeded cells in the chamber for 1 hour in 100 nM MLN8237 (Selleck Chemicals) prior to FRET imaging.

### Live cell imaging

For live imaging of S2 cells, cells were plated onto dishes with coverslip bottoms (MatTek Corporation, Ashland, MA, USA) previously treated with 0.2 mg/mL ConA (Sigma-Aldrich). Images were obtained at 30s intervals up to a maximum of 90min. All images were acquired at 25°C with either a spinning disc confocal system (Revolution; Andor) equipped with an electron multiplying charge-coupled device camera (iXonEM+; Andor) and a CSU-22 unit (Yokogawa) based on an inverted microscope (IX81; Olympus) driven by iQ software (Andor) or a Nikon ECLIPSE Ti2 (Nikon, Japan) equipped with Crest X-Light V3 spinning disk confocal system (Crest CrestOptics, Italy) and Kinetix 25 (Photometrics) camera driven by NIS elements 5.5 (Nikon, Japan) software. The microscopes are served with two laser lines, 488 and 561 nm, for the excitation of EGFP/GFP and mCherry, respectively. Time-lapse imaging of z stacks with 0.8 μm steps covering the entire volume of the cell was collected, and image sequence analysis and video assembly were done with Fiji software (https://fiji.sc/). The time each polar chromosome spent closer to the spindle pole then to the metaphase plate was quantified.

### Immunofluorescence imaging and analysis

For immunofluorescence analysis of kinetochore-microtubule attachments in S2 cells, 10^5^ cells were seeded onto coverslips previously treated with 0.2 mg/mL ConA (Sigma-Aldrich). For detection of calcium-stable k-fibers, cells were processed for fixation and immunostaining as previously described [35]. For immunofluorescence analysis of AAK^T311Ph^ levels in correlation to the distance between the centrosome and its closest kinetochore in S2 cells, 10^5^ cells were seeded onto coverslips previously treated with 0.2 mg/mL ConA (Sigma-Aldrich), processed for fixation in 4% paraformaldehyde in PBS for 12 min and further extracted for 8 min with 0.1% Triton X-100 in PBS, followed by immunostaining. For immunofluorescence analysis of AAK^T311Ph^ and ABK^T209Ph^ levels, 10^5^ cells were centrifuged onto slides for 5 min, at 239 g, and processed for fixation in 4% paraformaldehyde in PBS for 12 min and further extracted for 8 min with 0.1% Triton X-100 in PBS, followed by immunostaining. Immunostaining was performed as previously described [23]. Images were collected in Leica Scanning Confocal SP8 inverted microscope (Leica Microsystems). For immunofluorescence analysis of human cells, cells were grown on 12 mm round cover glasses (Menzel Glaser) and fixed by ice-cold methanol at -20°C for 4 minutes. DAPI was used at 1 μg/mL (Sigma) for DNA counterstaining. For quantification of centrosome to kinetochore distance, as well as for AAK and HEC1 phosphorylation status, images were acquired using a Plan-Apochromat DIC 63x/1.4 NA oil objective mounted on an inverted Zeiss Axio Observer Z1 microscope (Marianas Imaging Workstation from 3i-Intelligent Imaging and Innovations Inc.) equipped with a Hamamatsu ORCA-Flash4.0 v2 sCMOS Camera (Hamamatsu Photonics). Representative images were acquired using LSM700 confocal microscope (Carl Zeiss Inc.) mounted on a Zeiss-Axio imager Z1 equipped with alpha-Plan-Apochromat 100x/1.46 oil DIC M27 and plan-apochromat 63×/1.40 oil DIC M27 objective (Carl Zeiss Inc.) and Zen 2010 B SP1 software (Carl Zeiss, Inc.).

In S2 cells, the distance (in 3D) between the centrosome and its closest kinetochore was calculated using the measured values of the distance in the XY plane and the distance in the Z-axis (number of slices x Z-step). For immunofluorescence quantification, the mean fluorescence intensity of kinetochore or centrosome proteins was measured within a specific predefined region of interest (ROI) where individual kinetochores or centrosomes could fit. The kinetochores were identified based on their constitutive marker CID. The centrosomes were identified based on either their constitutive marker DPLP or, when stated, the transfected FLAG-AAK or EGFP-AAK versions. After background subtraction, identified as a region in the cell with no kinetochores or centrosomes, the intensities of the proteins of interest were determined relative to the respective constitutive markers, unless stated otherwise. Control values were averaged and used for normalization of values determined in the different biological conditions tested.

In human cells, kinetochore to centrosome distance and HEC1^S69Ph^ signal quantification on kinetochores in fixed cells were both measured using a custom routine written in MATLAB 8.2 (Mathworks). For analysis of kinetochore to centrosome distance, Pericentrin signal at centrosomes and CENP-C signal at kinetochores were used to determine the three-dimensional distance between the midpoint between the centrioles and the centroids of a kinetochore. For HEC1^S69Ph^ quantification on kinetochores, intensities were determined by quantification of pixel gray levels of the focused z plane within a ROI. Background fluorescence was measured outside the ROI and subtracted. For quantifications at kinetochores, the ROI encompassed a single kinetochore measuring the signal intensity of HEC1^S69Ph^ at the kinetochores. HEC1^S69Ph^ intensity was quantified relative to CENP-C fluorescence. HEC1^S69Ph^/CENP-C intensity ratios were subsequently normalized to control median value. At least 200 kinetochores from 10 different cells in each of three independent experiments were measured in both quantifications.

For AAK^T288Ph^ quantification at centrosomes, intensities were determined by measuring the signal intensity of AAK^T288Ph^ and total AAK in ImageJ software. Protein signal was quantified by drawing a circle closely along the total AAK staining at centrosomes and using the same ROI for AAK^T288Ph^ quantification. The ROI were adjusted in every cell to accurately match the total AAK staining. AAK^T288Ph^ intensity was quantified relative to total AAK fluorescence. AAK^T288Ph^/AAK intensity ratios were subsequently normalized to control median value. At least 25 cells in each of three independent experiments were used for the quantifications.

### Antibodies

The primary antibodies used for immunofluorescence in S2 cells were mouse anti-α-tubulin B512 (Sigma-Aldrich) used at 1:5000; rat anti-CID (Rat4) used at 1:500; chicken anti-GFP (#ab13970, Abcam, Cambridge, UK) used at 1:2,000; rabbit anti-CENP-C (Rb1, [36]; a gift from Christian Lehner, University of Zurich, Switzerland) used at 1:3000; guinea pig anti-MPS1 (Gp15, [37]; RRID:AB_2567774; a gift from Scott Hawley, The Stowers Institute for Medical Research, USA) used at 1:250; rabbit anti-phosphorylated Thr232-Aurora B (Rockland, Lim-erik, PA) used at 1:1000; rabbit anti-phosphorylated Thr288-Aurora A (C39D8, mAb #3079, Cell Signaling Technology, Danvers, MA, USA) used at 1:500; chicken anti-DPLP ([38]; a gift from Mónica Bettencourt-Dias, Instituto Gulbenkian de Ciencia, Portugal) used at 1:2000; mouse anti-FLAG (F1804, Sigma-Aldrich) used at 1:500. Alexa secondary antibodies were used according to the manufacturer’s instructions.

The primary antibodies used for immunofluorescence in human cells were guinea pig anti-CENP-C (PD030, MBL) used at 1:1000, mouse anti-Pericentrin (ab28144, Abcam) used at 1:1000, mouse anti-Aurora A (ab13824, Abcam) used at 1:200, rabbit anti-phosphorylated Thr288 Aurora A (AAK^T288Ph^) (C39D8 #3079, Cell Signalling) used at 1:1600 and rabbit anti-phosphorylated Ser69-HEC1 (HEC1^S69Ph^, [28]; a gift from Jennifer DeLuca, Colorado State University, USA) used at 1:3000. Goat anti-Mouse IgG (H + L) Highly Cross-Adsorbed Secondary Antibody, Alexa Fluor 568; Goat anti-Rabbit IgG (H + L) Highly Cross-Adsorbed Secondary Antibody, Alexa Fluor 488; and Goat anti-Guinea Pig IgG (H + L) Highly Cross-Adsorbed Secondary Antibody, Alexa Fluor 645 (Invitrogen) secondary antibodies were used at 1:1000.

The primary antibodies used for Western blotting analysis were rabbit anti-phosphorylated Thr676-MPS1 (MPS1^T676Ph^, [39]; a gift from Geert Kops, Oncode Institute, Hubrecht Institute, The Netherlands) used at 1:2000; rabbit anti-phosphorylated Thr288-Aurora A (AAK^T288Ph^, C39D8, mAb #3079, Cell Signaling Technology, Danvers, MA, USA) used at 1:1000.

### Expression and purification of recombinant proteins

For the mass spectrometry analysis, the catalytic domain of either human or *Drosophila* AAK wild-type or kinase-dead (AAK^K162I^ in human or AAK^K183I^ in *Drosophila*) were fused with N-terminal to MBP. To generate these constructs, human and *Drosophila* AAK cDNAs were used as template to amplify the regions corresponding to the amino acids 126-403 and 144-411, respectively. The PCR products were introduced in pMal-c2 (New England Biolabs) expression vector through FastCloning [34]. Recombinant vectors were subsequently transformed into BL21-star competent cells, and expression was induced overnight at 15°C by the addition of 0.05 mM IPTG. Lysates were sonicated and clarified by centrifugation at 4°C. The recombinant proteins were then incubated with MBP amylose magnetic beads (New England Biolabs) for 1 h at 4°C and washed with 200 mM NaCl, 20 mM Tris–HCl, 1 mM EDTA, 1 mM DTT, pH 7.4 and eluted in Column Buffer supplemented with 10 mM Maltose. The same protocol was followed for expression and purification of MBP protein alone.

### *In vitro* kinase assay and mass-spectrometry analysis

For in vitro kinase assays, recombinant human or *Drosophila* MBP-fused catalytic domain of wild-type or kinase-dead AAK, or 70nM of N-terminal His6-tagged, recombinant full-length human active AAK (#14-511 Merck Millipore, Germany) were incubated either in the presence or the absence of 14nM of N-terminal GST-fused active HsMPS1/TTK (SignalChem, Richmond, Canada) at 30°C for 30 min in a total volume of 30 μL kinase reaction buffer (5 mM MOPS, pH 7.2, 2.5 mM b-glycerol-phosphate, 5 mM MgCl_2_, 1 mM EGTA, 0.4 mM EDTA, 0.25 mM DTT, 0.1 mM ATP. For inhibition of HsMPS1/TTK activity, Compound 5 (Cpd-5) ([22]; gift from René Medema, Oncode Institute, The Netherlands Cancer Institute, The Netherlands) was added to the reaction at a final concentration of 100 nM. For AAK inhibition, MLN8237 (Selleck Chemicals) was added to the reaction at a final concentration of 100 nM. For Western blot analysis of AAK phosphorylation levels, reactions were stopped by addition of Laemmli sample buffer (4% SDS, 10% mercaptoetanol, 0.125 M Tris-HCl, 20% glycerol, 0.004% bromophenol blue) and heated for 5 min at 95°C. Peptides were resolved in an 10% SDS-PAGE. The activating autophosphorylation on T676 human MPS1/TTK and the activating autophosphorylation on T288 human AAK were detected by western blotting with phospho-specific antibodies. Total protein levels were detected by silver staining of the gel.

For identification of phosphorylated residues, the reaction was stopped by addition of 6 M urea and analyzed by liquid chromatography (LC) coupled to a mass spectrometer (LC/MS-MS). Samples were digested with LysC/Trypsin and prepared for LC-MS/MS analysis as previously described [40]. Peptides (100 ng) were separated on an EASY-nLC 1000 HPLC system (Thermo Fisher Scientific) with a 45 min gradient from 5–60% acetonitrile with 0.1% formic acid and directly sprayed via a nano-electrospray source in an Orbitrap Exploris^TM^ 480 (Thermo Fisher Scientific). The Q Exactive was operated in data-dependent mode, acquiring one survey scan and subsequently 15 MS/MS scans [41]. Resulting raw files were processed with MaxQuant software (v.16.14) using a reduced database containing the proteins of interest and a database of common contaminations. Oxidation (M) and phosphorylation (STY) were given as variable modifications and carbamidomethylation (C) as fixed modification [42]. A false discovery rate cut-off of 1% was applied at the peptide and protein levels and as well on phosphorylated peptides. Samples were run at least twice as technical replicates.

### Western blotting and gel staining

Coomassie staining of proteins resolved by SDS–PAGE was first conducted to determine comparable amounts of recombinant protein in solution to be used in the kinase assays. For Western blotting analysis, resolved proteins were transferred to a nitrocellulose membrane, using the iBlot Dry Blotting System (Thermo Fisher Scientific) according to the manufacturer’s instructions. Membranes were incubated for at least 1 h at room temperature in blocking solution (5% powder milk in PBS1×, 0.05% Tween 20). All primary and secondary antibodies were diluted in the blocking solution. Membranes were incubated with primary antibody solutions overnight at 4°C under constant stirring and washed three times in PBS1×, 0.05% Tween 20 for 10 min each.

Then, membranes were incubated with secondary antibody solutions for 1 h at room temperature under constant stirring. Secondary antibodies were conjugated to horseradish peroxidase (Jackson Immuno Research, UK). The blots were visualized by ECL detection and exposure to X-ray films (Fuji Medical X-Ray Film). Silver staining was conducted by gel fixation (in 30% ethanol, 10% acetic acid solution) for 2 h, followed by washing in 20% ethanol solution for 20 min and washing in water for 10 min. Then, the gel was incubated for 1min in 0,2g/L sodium thiosulfate, followed by a quick rinse with water and a 30 min incubation with 2g/L silver nitrate at 4°C. After a quick rinse with water, the gel was incubated with developer solution (10mg/L sodium thiosulfate, 0,025% formaldehyde, 30g/L sodium carbonate). When the adequate degree of staining was achieved, the development was stopped by adding a 50g/L Tris-base, 2,5% acetic acid solution.

### AlphaFold 3 predictions

AlphaFold 3 [43] was used to model the structure of *Drosophila melanogaster* AAK and the impact of T360 phosphorylation and T360A and T360D mutations. Models were obtained by inputting the kinase domain sequence (154-406 amino acids) of *Drosophila* AAK (Q9VGF9) as a single copy ‘Protein’ together with a single ADP ‘Ligand’ and two Mg2+ ‘Ion’ copies. Similarly, the mutants were obtained in the same conditions, but with either Alanine (A) or Aspartic acid (D) replacing Threonine (T) in amino acid position 360. For the Phospho-threonine (pT) model, the appropriate PTM was added onto the original sequence. For each of the five output models, ClashScore analysis and T-loop Confidence scores were assessed in SwissModel to determine the best model for each prediction. Confidence scores can be found in Figure S2H. All images of structures were processed on RCSB PBD - 3D view, where alignments are overall cartoon chain superpositions with ball and stick highlights on aa360, aa317, aa311 and molecular surface of the structure. The color coding was set as cyan blue for AAK^WT^, yellow for AAK^T360A^, red for AAK^T360D^ and orange for AAK^pT360^.

### Statistical analysis

All statistical analysis was performed with GraphPad Prism V10.0 (GraphPad Software, Inc.). Values were considered statistically different whenever P < 0.05. P values: ns, not significant; *< 0.05; **< 0.01; ***< 0.001; ****< 0.0001. *Drosophila* cells and western blot data are presented as median ± interquartile range or mean ± SD. Statistical analysis were performed using a Kruskal–Wallis test for multiple comparisons. Human cells data points were assessed for normality using the D’Agostino & Pearson test. Based on the results, statistical significance was evaluated either by Student’s t-test (unpaired, two-tailed for normally distributed data) or the Mann-Whitney U-test (unpaired, two-tailed for non-normally distributed data). Variances were compared using the F-test, and Welch’s correction was applied when variances were unequal. Detailed information on statistical significance for each condition can be found in the figures and their legends.

## REFERENCES

[1] Jelluma N, Brenkman AB, van den Broek NJ, Cruijsen CW, van Osch MH, Lens SM, Medema RH, Kops GJ. Mps1 phosphorylates Borealin to control Aurora B activity and chromosome alignment. Cell. 2008 Jan 25;132(2):233–46. doi: 10.1016/j.cell.2007.11.046. PMID: 18243099.

[2] Kwiatkowski N, Jelluma N, Filippakopoulos P, Soundararajan M, Manak MS, Kwon M, Choi HG, Sim T, Deveraux QL, Rottmann S, Pellman D, Shah JV, Kops GJ, Knapp S, Gray NS. Small-molecule kinase inhibitors provide insight into Mps1 cell cycle function. Nat Chem Biol. 2010 May;6(5):359–68. doi: 10.1038/nchembio.345. Epub 2010 Apr 11. PMID: 20383151; PMCID: PMC2857554.

[3] van der Waal MS, Saurin AT, Vromans MJ, Vleugel M, Wurzenberger C, Gerlich DW, Medema RH, Kops GJ, Lens SM. Mps1 promotes rapid centromere accumulation of Aurora B. EMBO Rep. 2012 Sep;13(9):847–54. doi: 10.1038/embor.2012.93. Epub 2012 Jun 26. PMID: 22732840; PMCID: PMC3432816.

[4] Santaguida S, Tighe A, D’Alise AM, Taylor SS, Musacchio A. Dissecting the role of MPS1 in chromosome biorientation and the spindle checkpoint through the small molecule inhibitor reversine. J Cell Biol. 2010 Jul 12;190(1):73–87. doi: 10.1083/jcb.201001036. PMID: 20624901; PMCID: PMC2911657.

[5] Hewitt L, Tighe A, Santaguida S, White AM, Jones CD, Musacchio A, Green S, Taylor SS. Sustained Mps1 activity is required in mitosis to recruit O-Mad2 to the Mad1-C-Mad2 core complex. J Cell Biol. 2010 Jul 12;190(1):25–34. doi: 10.1083/jcb.201002133. PMID: 20624899; PMCID: PMC2911659.

[6] Maciejowski J, George KA, Terret ME, Zhang C, Shokat KM, Jallepalli PV. Mps1 directs the assembly of Cdc20 inhibitory complexes during interphase and mitosis to control M phase timing and spindle checkpoint signaling. J Cell Biol. 2010 Jul 12;190(1):89–100. doi: 10.1083/jcb.201001050. PMID: 20624902; PMCID: PMC2911671.

[7] Hayward D, Roberts E, Gruneberg U. MPS1 localizes to end-on microtubule-attached kinetochores to promote microtubule release. Curr Biol. 2022 Dec 5;32(23):5200–5208.e8. doi: 10.1016/j.cub.2022.10.047. Epub 2022 Nov 16. PMID: 36395767.

[8] Maciejowski J, Drechsler H, Grundner-Culemann K, Ballister ER, Rodriguez-Rodriguez JA, Rodriguez-Bravo V, Jones MJK, Foley E, Lampson MA, Daub H, McAinsh AD, Jallepalli PV. Mps1 Regulates Kinetochore-Microtubule Attachment Stability via the Ska Complex to Ensure Error-Free Chromosome Segregation. Dev Cell. 2017 Apr 24;41(2):143–156.e6. doi: 10.1016/j.devcel.2017.03.025. PMID: 28441529; PMCID: PMC5477644.

[9] Sarangapani KK, Koch LB, Nelson CR, Asbury CL, Biggins S. Kinetochore-bound Mps1 regulates kinetochore-microtubule attachments via Ndc80 phosphorylation. J Cell Biol. 2021 Dec 6;220(12):e202106130. doi: 10.1083/jcb.202106130. Epub 2021 Oct 14. PMID: 34647959; PMCID: PMC8641409.

[10] Chmátal L, Yang K, Schultz RM, Lampson MA. Spatial Regulation of Kinetochore Microtubule Attachments by Destabilization at Spindle Poles in Meiosis I. Curr Biol. 2015 Jul 20;25(14):1835–41. doi: 10.1016/j.cub.2015.05.013. Epub 2015 Jul 9. PMID: 26166779; PMCID: PMC4519087.

[11] Ye AA, Deretic J, Hoel CM, Hinman AW, Cimini D, Welburn JP, Maresca TJ. Aurora A Kinase Contributes to a Pole-Based Error Correction Pathway. Curr Biol. 2015 Jul 20;25(14):1842–51. doi: 10.1016/j.cub.2015.06.021. Epub 2015 Jul 9. PMID: 26166783; PMCID: PMC4509859.

[12] Moura M, Osswald M, Leça N, Barbosa J, Pereira AJ, Maiato H, Sunkel CE, Conde C. Protein Phosphatase 1 inactivates Mps1 to ensure efficient Spindle Assembly Checkpoint silencing. Elife. 2017 May 2;6:e25366. doi: 10.7554/eLife.25366. PMID: 28463114; PMCID: PMC5433843.

[13] Dou Z, Liu X, Wang W, Zhu T, Wang X, Xu L, Abrieu A, Fu C, Hill DL, Yao X. Dynamic localization of Mps1 kinase to kinetochores is essential for accurate spindle microtubule attachment. Proc Natl Acad Sci U S A. 2015 Aug 18;112(33):E4546–55. doi: 10.1073/pnas.1508791112. Epub 2015 Aug 3. PMID: 26240331; PMCID: PMC4547264.

[14] Hiruma Y, Sacristan C, Pachis ST, Adamopoulos A, Kuijt T, Ubbink M, von Castelmur E, Perrakis A, Kops GJ. CELL DIVISION CYCLE. Competition between MPS1 and microtubules at kinetochores regulates spindle checkpoint signaling. Science. 2015 Jun 12;348(6240):1264–7. doi: 10.1126/science.aaa4055. Epub 2015 Jun 11. PMID: 26068855.

[15] Ji Z, Gao H, Yu H. CELL DIVISION CYCLE. Kinetochore attachment sensed by competitive Mps1 and microtubule binding to Ndc80C. Science. 2015 Jun 12;348(6240):1260–4. doi: 10.1126/science.aaa4029. PMID: 26068854.

[16] Parnell EJ, Jenson EE, Miller MP. A conserved site on Ndc80 complex facilitates dynamic recruitment of Mps1 to yeast kinetochores to promote accurate chromosome segregation. Curr Biol. 2024 May 16:S0960-9822(24)00528–1. doi: 10.1016/j.cub.2024.04.054. Epub ahead of print. PMID: 38776906.

[17] Pleuger R, Cozma C, Hohoff S, Denkhaus C, Dudziak A, Kaschani F, Kaiser M, Musacchio A, Vetter IR, Westermann S. Microtubule end-on attachment maturation regulates Mps1 association with its kinetochore receptor. Curr Biol. 2024 May 16:S0960-9822(24)00402–0. doi: 10.1016/j.cub.2024.03.062. Epub ahead of print. PMID: 38776902.

[18] Zahm JA, Harrison SC. A communication hub for phosphoregulation of kinetochore-microtubule attachment. Curr Biol. 2024 May 16:S0960-9822(24)00580–3. doi: 10.1016/j.cub.2024.04.067. Epub ahead of print. PMID: 38776904.

[19] Fuller BG, Lampson MA, Foley EA, Rosasco-Nitcher S, Le KV, Tobelmann P, Brautigan DL, Stukenberg PT, Kapoor TM. Midzone activation of aurora B in anaphase produces an intracellular phosphorylation gradient. Nature. 2008 Jun 19;453(7198):1132–6. doi: 10.1038/nature06923. Epub 2008 May 7. PMID: 18463638; PMCID: PMC2724008.

[20] Violin JD, Zhang J, Tsien RY, Newton AC. A genetically encoded fluorescent reporter reveals oscillatory phosphorylation by protein kinase C. J Cell Biol. 2003 Jun 9;161(5):899–909. doi: 10.1083/jcb.200302125. Epub 2003 Jun 2. PMID: 12782683; PMCID: PMC2172956.

[21] Feng Z, Caballe A, Wainman A, Johnson S, Haensele AFM, Cottee MA, Conduit PT, Lea SM, Raff JW. Structural Basis for Mitotic Centrosome Assembly in Flies. Cell. 2017 Jun 1;169(6):1078–1089.e13. doi: 10.1016/j.cell.2017.05.030. PMID: 28575671; PMCID: PMC5457487.

[22] Koch A, Maia A, Janssen A, Medema RH. Molecular basis underlying resistance to Mps1/TTK inhibitors. Oncogene. 2016 May 12;35(19):2518–28. doi: 10.1038/onc.2015.319. Epub 2015 Sep 14. PMID: 26364596; PMCID: PMC4867491.

[23] Conde C, Osswald M, Barbosa J, Moutinho-Santos T, Pinheiro D, Guimarães S, Matos I, Maiato H, Sunkel CE. Drosophila Polo regulates the spindle assembly checkpoint through Mps1-dependent BubR1 phosphorylation. EMBO J. 2013 Jun 12;32(12):1761–77. doi: 10.1038/emboj.2013.109. Epub 2013 May 17. PMID: 23685359; PMCID: PMC3680734.

[24] Lamb JR, Michaud WA, Sikorski RS, Hieter PA. Cdc16p, Cdc23p and Cdc27p form a complex essential for mitosis. EMBO J. 1994 Sep 15;13(18):4321–8. doi: 10.1002/j.1460-2075.1994.tb06752.x. PMID: 7925276; PMCID: PMC395359.

[25] Tugendreich S, Tomkiel J, Earnshaw W, Hieter P. CDC27Hs colocalizes with CDC16Hs to the centrosome and mitotic spindle and is essential for the metaphase to anaphase transition. Cell. 1995 Apr 21;81(2):261–8. doi: 10.1016/0092-8674(95)90336-4. PMID: 7736578.

[26] Huang JY, Raff JW. The dynamic localisation of the Drosophila APC/C: evidence for the existence of multiple complexes that perform distinct functions and are differentially localised. J Cell Sci. 2002 Jul 15;115(Pt 14):2847–56. doi: 10.1242/jcs.115.14.2847. PMID: 12082146.

[27] Barisic M, Aguiar P, Geley S, Maiato H. Kinetochore motors drive congression of peripheral polar chromosomes by overcoming random arm-ejection forces. Nat Cell Biol. 2014 Dec;16(12):1249–56. doi: 10.1038/ncb3060. Epub 2014 Nov 10. PMID: 25383660.

[28] DeLuca KF, Meppelink A, Broad AJ, Mick JE, Peersen OB, Pektas S, Lens SMA, DeLuca JG. Aurora A kinase phosphorylates Hec1 to regulate metaphase kinetochore-microtubule dynamics. J Cell Biol. 2018 Jan 2;217(1):163–177. doi: 10.1083/jcb.201707160. Epub 2017 Nov 29. PMID: 29187526; PMCID: PMC5748988.

[29] Ye AA, Maresca TJ. It’s all relative: Centromere-versus pole-based error correction. Cell Cycle. 2015;14(24):3777–8. doi: 10.1080/15384101.2015.1105701. PMID: 26697831; PMCID: PMC4825761.

[30] Eibes S, Rajendraprasad G, Guasch-Boldu C, Kubat M, Steblyanko Y, Barisic M. CENP-E activation by Aurora A and B controls kinetochore fibrous corona disassembly. Nat Commun. 2023 Sep 1;14(1):5317. doi: 10.1038/s41467-023-41091-2. PMID: 37658044; PMCID: PMC10474297.

[31] Sikirzhytski V, Renda F, Tikhonenko I, Magidson V, McEwen BF, Khodjakov A. Microtubules assemble near most kinetochores during early prometaphase in human cells. J Cell Biol. 2018 Aug 6;217(8):2647–2659. doi: 10.1083/jcb.201710094. Epub 2018 Jun 15. PMID: 29907657; PMCID: PMC6080938.

[32] Chakraborty M, Tarasovetc EV, Zaytsev AV, Godzi M, Figueiredo AC, Ataullakhanov FI, Grishchuk EL. Microtubule end conversion mediated by motors and diffusing proteins with no intrinsic microtubule end-binding activity. Nat Commun. 2019 Apr 11;10(1):1673. doi: 10.1038/s41467-019-09411-7. PMID: 30975984; PMCID: PMC6459870.

[33] Renda F, Miles C, Tikhonenko I, Fisher R, Carlini L, Kapoor TM, Mogilner A, Khodjakov A. Non-centrosomal microtubules at kinetochores promote rapid chromosome biorientation during mitosis in human cells. Curr Biol. 2022 Mar 14;32(5):1049–1063.e4. doi: 10.1016/j.cub.2022.01.013. Epub 2022 Feb 1. PMID: 35108523; PMCID: PMC8930511.

[34] Li C, Wen A, Shen B, Lu J, Huang Y, Chang Y. FastCloning: a highly simplified, purification-free, sequence- and ligation-independent PCR cloning method. BMC Biotechnol. 2011 Oct 12;11:92. doi: 10.1186/1472-6750-11-92. PMID: 21992524; PMCID: PMC3207894.

[35] Kapoor TM, Mayer TU, Coughlin ML, Mitchison TJ. Probing spindle assembly mechanisms with monastrol, a small molecule inhibitor of the mitotic kinesin, Eg5. J Cell Biol. 2000 Sep 4;150(5):975–88. doi: 10.1083/jcb.150.5.975. PMID: 10973989; PMCID: PMC2175262.

[36] Heeger S, Leismann O, Schittenhelm R, Schraidt O, Heidmann S, Lehner CF. Genetic interactions of separase regulatory subunits reveal the diverged Drosophila Cenp-C homolog. Genes Dev. 2005 Sep 1;19(17):2041–53. doi: 10.1101/gad.347805. PMID: 16140985; PMCID: PMC1199574.

[37] Gilliland WD, Hughes SE, Cotitta JL, Takeo S, Xiang Y, Hawley RS. The multiple roles of mps1 in Drosophila female meiosis. PLoS Genet. 2007 Jul;3(7):e113. doi: 10.1371/journal.pgen.0030113. PMID: 17630834; PMCID: PMC1914070.

[38] Rodrigues-Martins A, Riparbelli M, Callaini G, Glover DM, Bettencourt-Dias M. Revisiting the role of the mother centriole in centriole biogenesis. Science. 2007 May 18;316(5827):1046-50. doi: 10.1126/science.1142950. Epub 2007 Apr 26. PMID: 17463247.

[39] Jelluma N, Brenkman AB, McLeod I, Yates JR 3rd, Cleveland DW, Medema RH, Kops GJ. Chromosomal instability by inefficient Mps1 auto-activation due to a weakened mitotic checkpoint and lagging chromosomes. PLoS One. 2008 Jun 11;3(6):e2415. doi: 10.1371/journal.pone.0002415. PMID: 18545697; PMCID: PMC2408436.

[40] Rappsilber J, Mann M, Ishihama Y. Protocol for micro-purification, enrichment, pre-fractionation and storage of peptides for proteomics using StageTips. Nat Protoc. 2007;2(8):1896–906. doi: 10.1038/nprot.2007.261. PMID: 17703201.

[41] Olsen JV, Macek B, Lange O, Makarov A, Horning S, Mann M. Higher-energy C-trap dissociation for peptide modification analysis. Nat Methods. 2007 Sep;4(9):709–12. doi: 10.1038/nmeth1060. Epub 2007 Aug 26. PMID: 17721543.

[42] Cox J, Mann M. MaxQuant enables high peptide identification rates, individualized p.p.b.-range mass accuracies and proteome-wide protein quantification. Nat Biotechnol. 2008 Dec;26(12):1367–72. doi: 10.1038/nbt.1511. Epub 2008 Nov 30. PMID: 19029910.

[43] Abramson J, Adler J, Dunger J, Evans R, Green T, Pritzel A, Ronneberger O, Willmore L, Ballard AJ, Bambrick J, Bodenstein SW, Evans DA, Hung CC, O’Neill M, Reiman D, Tunyasuvunakool K, Wu Z, Žemgulytė A, Arvaniti E, Beattie C, Bertolli O, Bridgland A, Cherepanov A, Congreve M, Cowen-Rivers AI, Cowie A, Figurnov M, Fuchs FB, Gladman H, Jain R, Khan YA, Low CMR, Perlin K, Potapenko A, Savy P, Singh S, Stecula A, Thillaisundaram A, Tong C, Yakneen S, Zhong ED, Zielinski M, Žídek A, Bapst V, Kohli P, Jaderberg M, Hassabis D, Jumper JM. Accurate structure prediction of biomolecular interactions with AlphaFold 3. Nature. 2024 May 8. doi: 10.1038/s41586-024-07487-w. Epub ahead of print. PMID: 38718835.

