## Supplementary figures and images for "Proximity-based activation of AURORA A by MPS1 potentiates error correction"

### Supplemental Figure 1

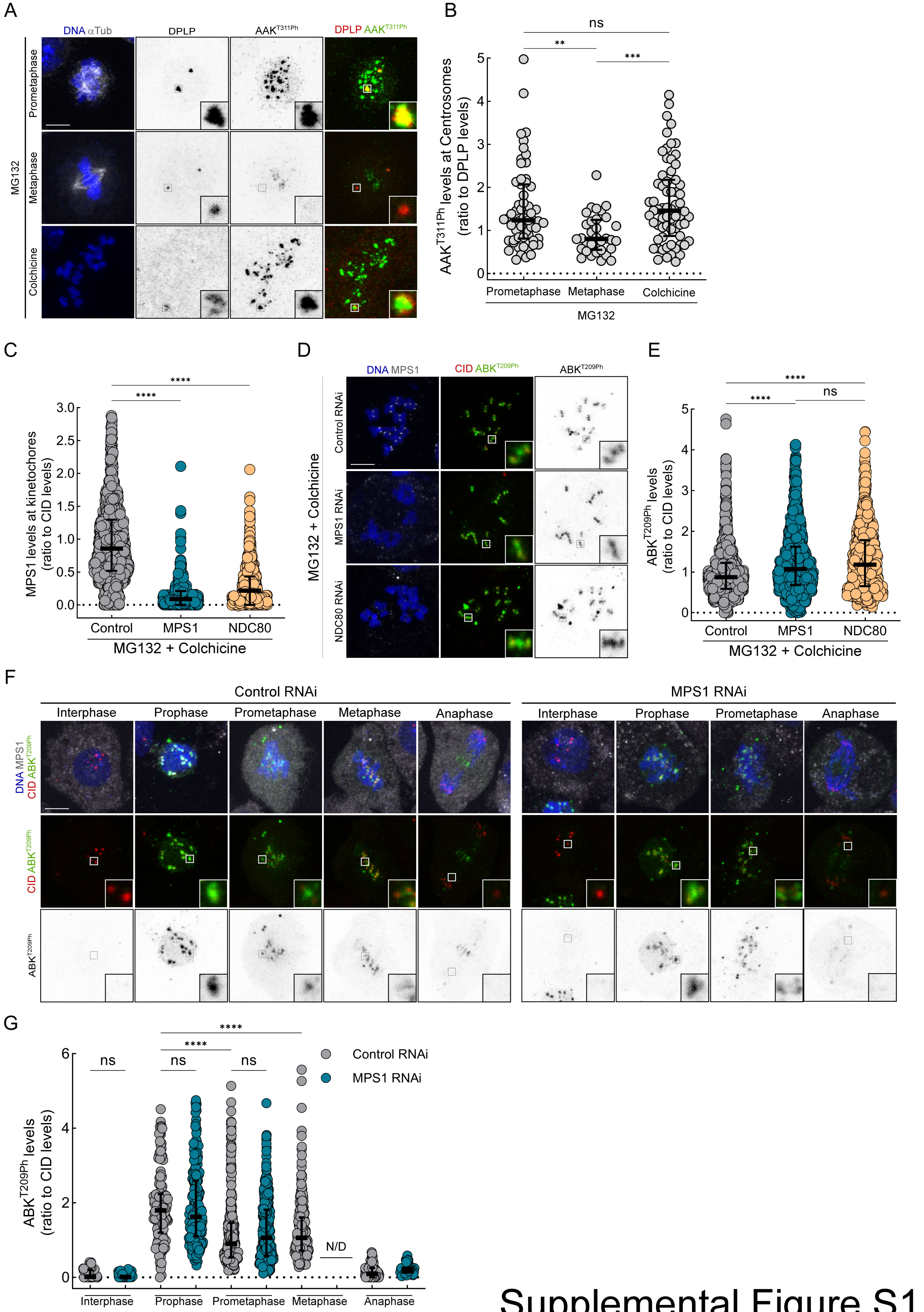


#### Supplemental Figure S1

### Supplemental Figure 3

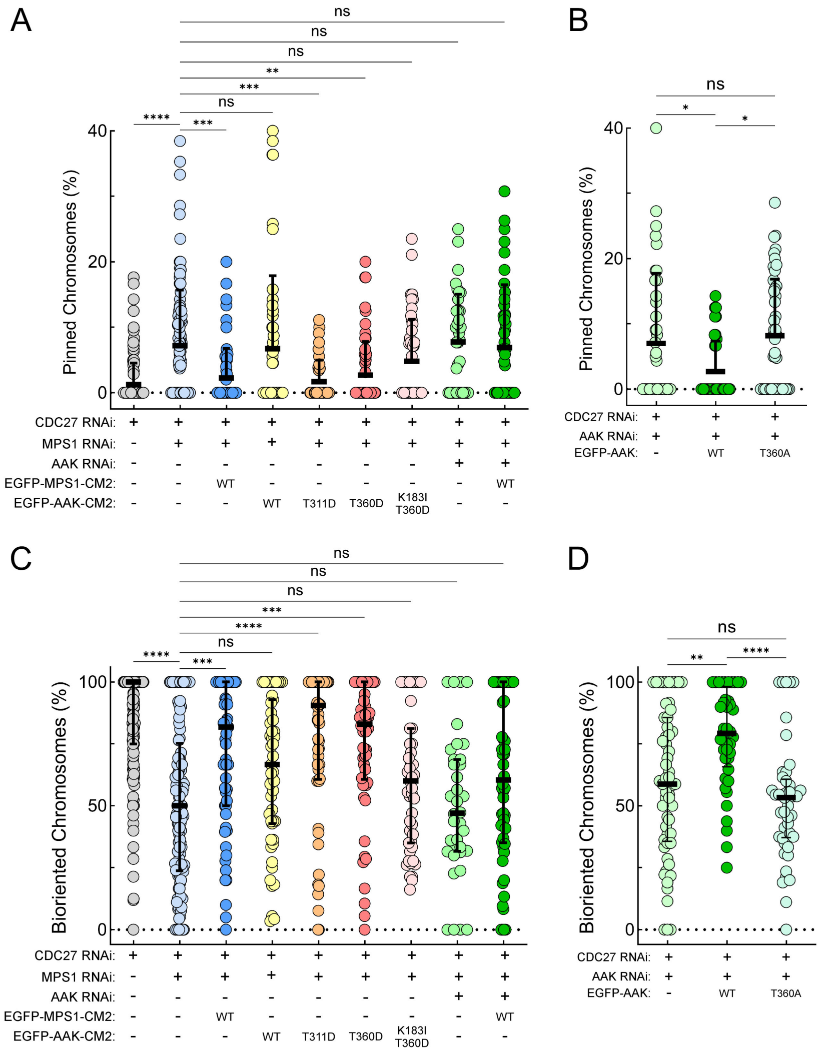


#### Supplemental Figure S3

### Supplemental Figure 4

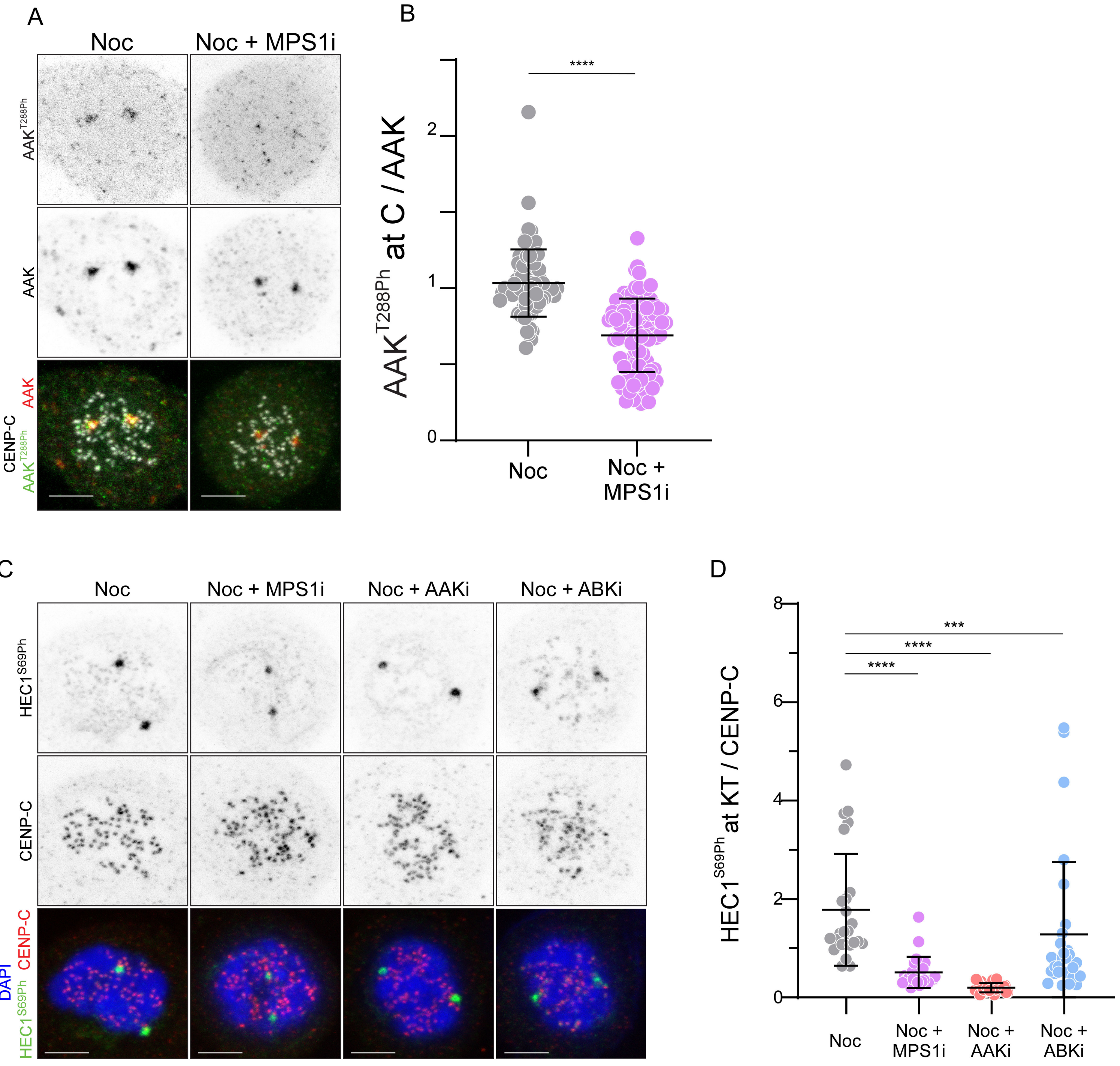


Supplemental Figure S4
