## Supplemental Figure 2 for "Proximity-based activation of AURORA A by MPS1 potentiates error correction"

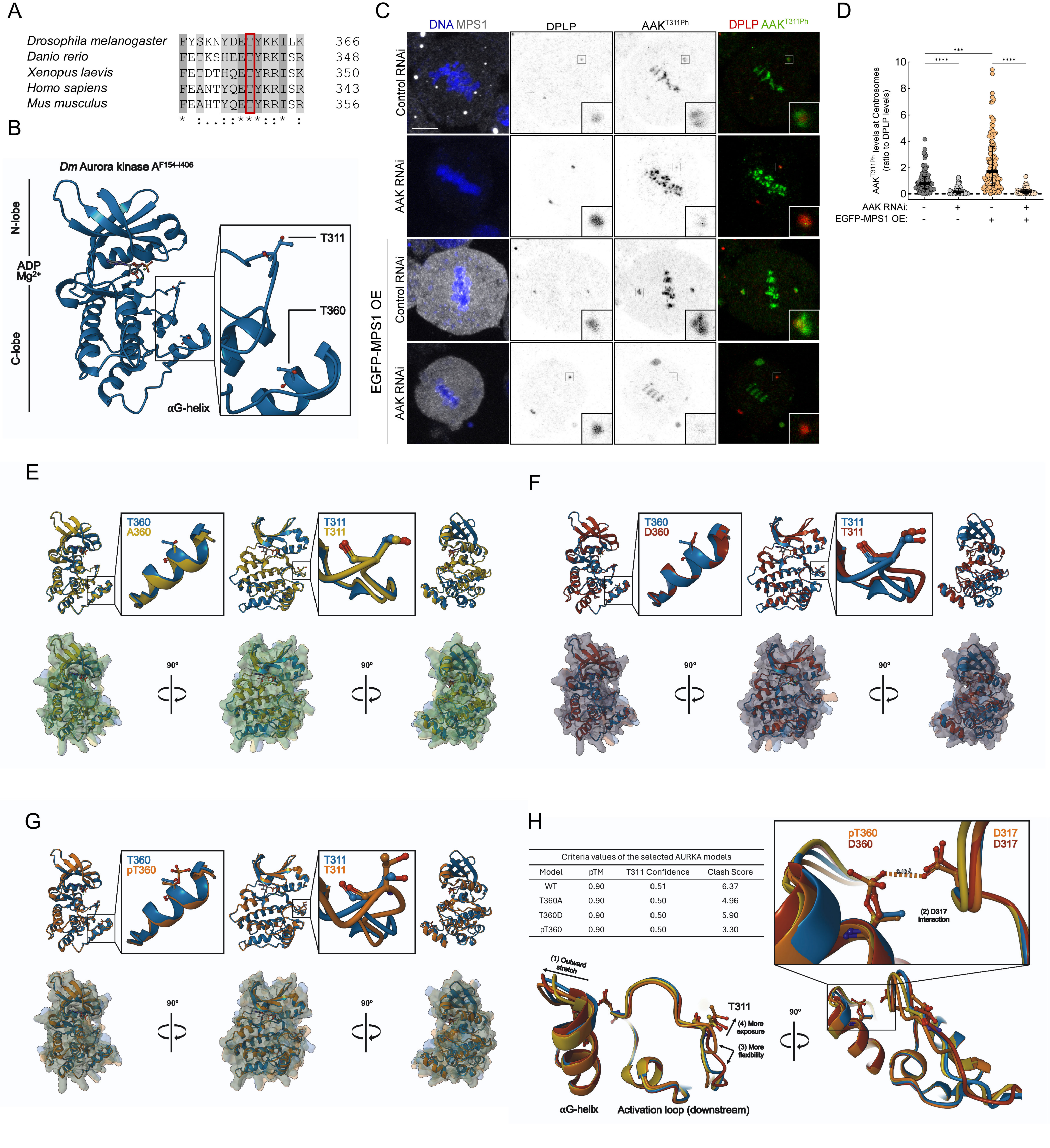


Criteria values of the selected AURKA models

| Model | pTM | T311 Confidence | Clash Score |
|---|---|---|---|
| WT | 0.90 | 0.51 | 6.37 |
| T360A | 0.90 | 0.50 | 4.96 |
| T360D | 0.90 | 0.50 | 5.90 |
| pT360 | 0.90 | 0.50 | 3.30 |

### Supplemental Figure S2
